# Sustained Delivery of GLP-1 Receptor Agonists from Injectable Biomimetic Hydrogels Improves Treatment of Diabetes

**DOI:** 10.1101/2023.01.28.526057

**Authors:** Andrea I. d’Aquino, Caitlin L. Maikawa, Leslee T. Nguyen, Katie Lu, Ian A. Hall, Carolyn K. Jons, Catherine M. Kasse, Jerry Yan, Alexander N. Prossnitz, Enmian Chang, Sam W. Baker, Lars Hovgaard, Dorte B. Steensgaard, Hanne B. Andersen, Lotte Simonsen, Eric A. Appel

## Abstract

Glucagon-like peptide-1 (GLP-1) is an incretin hormone and neurotransmitter secreted from intestinal L-cells in response to nutrients to stimulate insulin and block glucagon secretion in a glucose-dependent manner. GLP-1 in itself is rapidly degraded, but long-acting GLP-1 receptor agonists (GLP-1 RAs) have become central in the treatment of T2D because of the beneficial effects extending also beyond glucose control. Currently, these therapeutics must be injected either daily or weekly or taken daily orally, leaving room for technological innovations that enable less frequent administrations, which will reduce patient burden and increase patient compliance. An ideal GLP-1 RA drug product would provide continuous therapy for upwards of four months from a single administration to match the cadence with which T2D patients typically visit their physician. In this work, we leveraged an injectable hydrogel depot technology to develop a long-acting GLP-1 RA drug product. By modulating the hydrogel properties to tune GLP-1 RA retention within the hydrogel depot, we engineered formulations capable of months-long GLP-1 RA delivery. Using a rat model of T2D, we confirmed that a single injection of hydrogel-based therapies exhibits sustained exposure of GLP-1 RA over 42 days, corresponding to a once-every four month therapy in humans. Moreover, these hydrogel therapies maintained optimal management of blood glucose and weight comparable to daily injections of a leading GLP-1 RA drug molecule. The pharmacokinetics and pharmacodynamics of these hydrogel-based long-acting GLP-1 RA treatments are promising for development of novel therapies reducing treatment burden for more effective management of T2D.

**Progress and Potential:** While insufficient access to quality healthcare is problematic for consistent management of Type II diabetes (T2D), poor adherence to burdensome treatment regimens is one of the greatest challenges for disease management. Glucagon-like peptide 1 (GLP1) drugs have become central to the treatment of T2D due to their many beneficial effects beyond improving glucose control. Unfortunately, while optimization of GLP1 drugs has reduced treatment frequency from daily to weekly, significant patient burden still leads to poor patience compliance. In this work we developed an injectable hydrogel technology to enable GLP1 drugs only requiring administration once every four months. We showed in a rat model of T2D that one injection of a hydrogel-based therapy improves management of blood glucose and weight when compared with daily injections of the leading drug used clinically. These hydrogel-based GLP1 treatments are promising for reducing treatment burden and more effectively managing T2D.

**Future Impact:** A GLP-1-based drug product providing four months of continuous therapy per administration could be transformational for the management of Type II diabetes (T2D). One of the most challenging aspects of diabetes management with GLP-1 mimics is maintenance of consistent levels of the drugs in the body, which is complicated by poor patient compliance on account of the high frequency of dosing required for current treatments. By leveraging a unique sustained release hydrogel depot technology we develop a months-long GLP-1 drug product candidate that has the potential to reduce patient burden and improving diabetes management. Overall, the hydrogel technology we describe here can dramatically reduce the frequency of therapeutic interventions, significantly increasing patient quality of life and reducing complications of diabetes management.

Our next steps will focus on optimization of the drug formulations in a swine model of T2D, which is the most advanced and translationally-relevant animal model for these types of therapeutics. The long-term vision for this work is to translate lead candidate drug products towards clinical evaluation, which will also require comprehensive safety evaluation in multiple species and manufacturing our these materials according to Good Manufacturing Practices. The months-long-acting GLP-1 drug product that will come from this work has the potential to afford thus far unrealized therapeutic impact for the hundreds of millions of people with diabetes worldwide.

## Main Text

Out of almost 500 million people living with diabetes or pre-diabetes worldwide, an estimated 130 million live in the US (*2, 3*). In the US alone, the annual spendings directely related to diabetes and pre-diabetes amounts to roughly $400 billion. This makes it the tenth most costly disease in the US (*2, 4*). Type 2 diabetes (T2D), which accounts for 90–95% of all diabetes cases, is a metabolic disorder characterized by insulin resistance, deterioration of pancreatic β-cell function, and impaired regulation of hepatic glucose production eventually leading to β-cell failure (*5–7*). Patients with poorly managed T2D are at risk of serious micro- and macrovascular complications, including cardiovascular disease, nephropathy, retinopathy, neuropathy and stroke. Besides insufficient access to care, poor adherence to treatment is a general problem (*1, 8*). The lack of adherence occurs primarily due to adverse effects, most commonly being hypoglycemia, weight gain or gastrointestinal side effects, and complex treatment regimens (*1, 8*). Treatment strategies that alleviate patient burden while providing optimal glycemic control would transform the treatment of T2D and improve treatment outcomes and patient life.

Glucagon-like peptide-1 receptor agonists (GLP-1 RAs) have become central to the treatment of T2D due to their beneficial effects that extend beyond improving glucose control (*9, 10*). GLP-1 is an incretin hormone secreted from intestinal L-cells in response to nutrients that lowers blood glucose by stimulating insulin and suppressing glucagon secretion in a glucose-dependent manner, reducing the risk of hypoglycemia (*11*). In addition, GLP-1 is also a neurotransmitter synthesized by preproglucagon neurons in the brain and acts via central pathways to lower energy intake through an effect on satiety, hunger and reward related measures (*12, 13*), leading to a lowering of body weight, which has prompted the approval of liraglutide and semaglutide for the treatment of obesity (*14–18*). Importantly, liraglutide is currently the only GLP-1 RA that is FDA approved for use in children ages 12-17 years with obesity (*19*). Long-acting GLP-1 RAs also reduce risk of cardiovascular disease (*20–22*), making them attractive treatment options for people who are at increased risk of these disorders. Importantly, it has been established that optimal therapeutic effect of GLP-1 is achieved with regimens that provide sustained plasma levels of active peptide (*9–11*). Such regimens include continuous subcutaneous infusion of GLP-1 (*12–14*), and once- or twice-daily or weekly injections of GLP-1 analogues with extended plasma stability (*14, 15*). Therefore, novel delivery technologies that provide sustained plasma levels of GLP-1 RAs is particularly important for treating diabetes.

Several strategies have been employed for stabilizing GLP-1 RAs and prolonging therapeutic efficacy, resulting in development of T2D drug products with treatment regimens ranging from twice daily to once weekly (twice daily and weekly exenatide; daily liraglutide; weekly dulaglutide; and weekly semaglutide) (*10*). Liraglutide and semaglutide are based on the native GLP-1 sequence, carry a single optimized fatty acid and linker modification enabling reversible binding to albumin to extend the circulating half-life while maintaining optimal potency. While liraglutide has a pharmacokinetic profile supporting once daily dosing, semaglutide has extended pharmacokinetics and once weekly dosing enabled by further improved metabolic stability and a di-carboxylic fatty acid sidechain facilitating a stronger binding to albumin (*23*). While reducing treatment frequency from daily to weekly is associated with improved patient adherence, there is still room for improvement to reduce treatment burden and improve patience compliance (Figure 1A) (*24–26*). To address this challenge, we sought to develop long-acting formulations of semaglutide and liraglutide providing continuous therapy for upwards of four months from a single administration to coincide with the typical cadence with which T2D patients visit their endocrinologist or primary care provider (Figure 1B and C) (*27*).

**Fig. 1.**
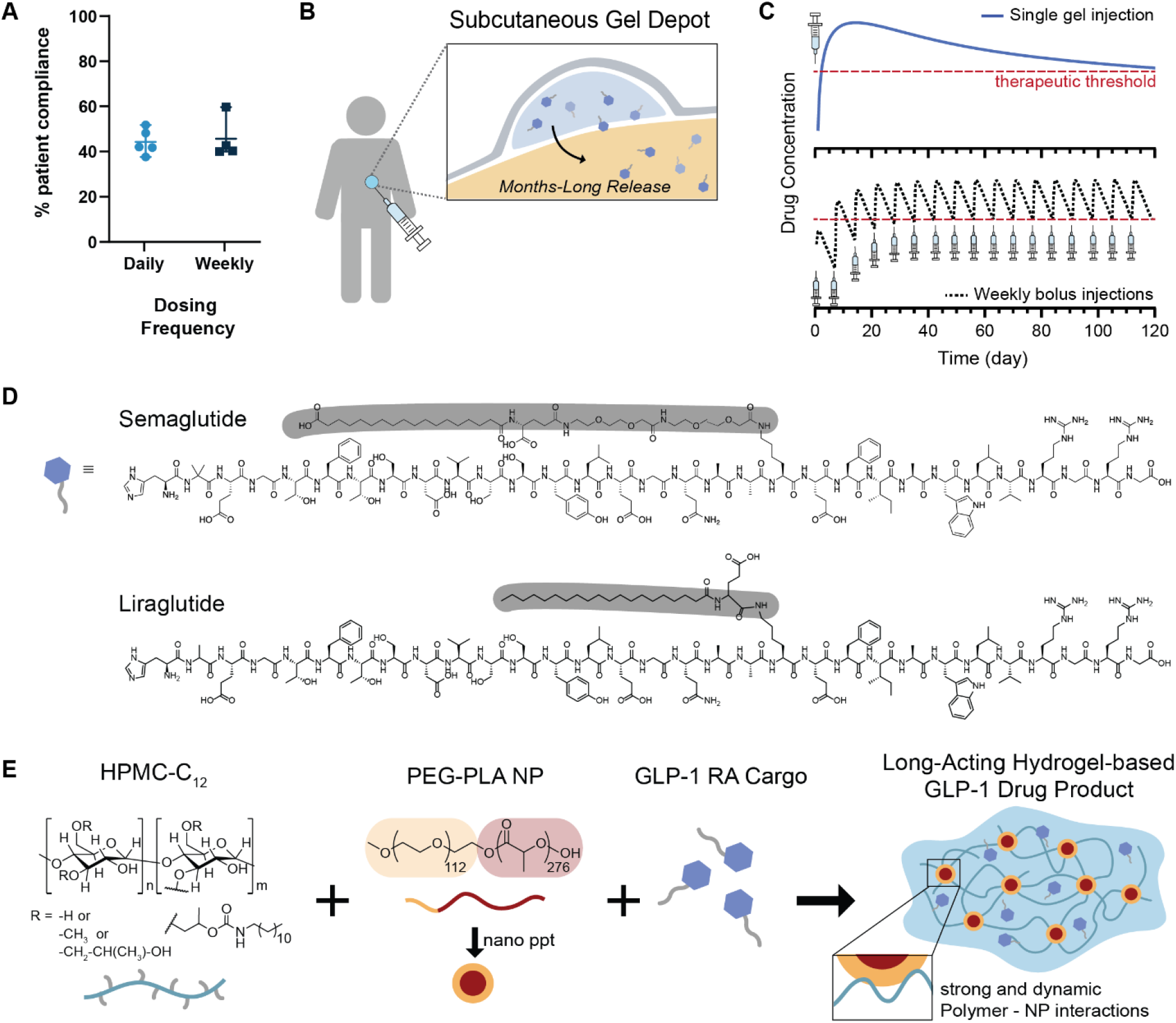
PNP hydrogels for the prolonged delivery of GLP-1 Ras. A) From literature reports it is clear that once weekly dosing frequency does not significantly improve patient compliance compared to a once daily dosing frequency (*1*). B) Localized depots form in the subcutaneous space immediately after subcutaneous injection, providing a tunable platform for sustained release of GLP-1-RA compounds. C) Clinical data showing the release profile of current GLP-1 treatments, where the black dotted line represents repeated weekly injections that patients take every week for four months to reach therapeutic concentrations of GLP-1. In contrast, the blue line represents the target delivery profile of a single PNP hydrogel depot injection that sustains release of GLP-1 for 120 days. Current state-of-the-art strategies require daily or weekly subcutaneous injections with significant ramp up time to achieve therapeutic concentrations. The red dotted line indicates the therapeutic threshold. D) Semaglutide and liraglutide are once-weekly and once-daily GLP-1 RA therapies, respectively, that were investigated in this study. E) PNP hydrogels prepared by mixing hydrophobically-modified HPMC with PEG-PLA nanoparticles enable facile encapsulation of GLP-1-receptor agonists with 100% efficiency.

In this work, we drew inspiration from long-acting drug products such as Lupron Depot® containing leuprorelin, which is an effective microparticle-based leuprorelin formulation for treatment of endometriosis developed for administration once every three months (*28*). Additionally, one-month, three-month, four-month and six-month microparticle-based leuprorelin depot formulations are widely used for treatment of advanced and metastatic prostate cancer (*29*). Similary, exenatide is commercially available in a microparticle-based extended-release formulation enabling once-weekly administration. While these microparticle-based drug delivery technologies have significantly improved the utility of exenatide as a pharmaceutical agent by enabling once weekly dosing, they have unfortunately required significant optimization to achieve appropriate drug-depot compatibility and release behavior, and only relatively hydrophobic peptides have demonstrated months-long release (*30*). For this reason, we sought to engineer an injectable hydrogel depot technology capable of months-long release of GLP-1 RAs. Hydrogels address some of the shortcomings of microparticle technologies as they maintain the native aqueous environment around encapsulated drugs and are therefore compatible with existing approved drug molecules developed for aqueous formulation. While many hydrogel-based depot technologies have been reported in the literature to show attractive local tolerance (*31*), they typically exhibit several critical shortcomings, including: (i) complicated manufacturing and poor formulation stability, (ii) challenging administration, (iii) burst release that can contribute to poor tolerability of the therapy, and (iv) insufficiently slow release to enable appropriately long-acting therapies. We therefore sought to leverage an injectable hydrogel platform generated through self-assembly of dynamic, entropically-driven supramolecular interactions between biodegradable nanoparticles (NPs) and hydrophobically-modified hydroxypropyl methylcellulose (HPMC) derivatives to develop a long-acting GLP-1 RA formulation (Figure 1E) (*32–39*). We have previously shown these polymer-nanoparticle (PNP) hydrogels to be capable of providing application-specific release of diverse biopharmaceuticals such as proteins, vaccines, and cells that is tunable over timeframes extending from days to upwards of six months (*33, 37, 40–44*).

Unlike traditional covalently crosslinked hydrogels, PNP hydrogels are formed through strong yet dynamic physical interactions. As a result, these materials address the shortcomings of other hydrogel-based depot technologies as they exhibit: (i) mild formulation requirements favorable for maintaining drug stability during manufacturing and storage, (ii) pronounced shear thinning properties enabling injection through clinically-relevant needles, (iii) rapid self-healing of hydrogel structure and depot formation mitigating burst release of the drug cargo, (iv) sufficiently high yield stress to form a robust depot that persists under the normal stresses of the subcutaneous space following administration, and (v) prolonged delivery of therapeutic cargo (*45*). These hydrogels are by design biodegradable and have been shown to be non-immunogenic in mice, rats, pigs and sheep, as well as not promoting immune responses to their payloads (*33–39, 41, 44*). Moreover, these gels can be formulated and stored in prefilled syringes as they maintain excellent stability under standard storage conditions (e.g., refrigeration) (*39*). Herein, we sought to optimize GLP-1 RA pharmacokinetics by engineering the drug encapsulation and release characteristics of these PNP hydrogels, thereby enabling the development of a months-long-acting GLP-1 RA drug product capable of reducing treatment burden to more effectively manage T2D.

## RESULTS

### Development of injectable hydrogels for sustained release of GLP-1 RAs

Liraglutide and semaglutide were considered excellent therapeutic cargo for sustained delivery from PNP hydrogel depot materials due to the hydrophobic fatty acid modifications (*23, 46, 47*). We hypothesized that these fatty acid moieties could non-covalently embed the biopharmaceutical within the hydrogel network with simple mixing during hydrogel fabrication (Figure 1D). This embedding is made possible by the fact that the poly(lactic acid) core of the PEG-PLA NPs within the hydrogel structure is hydrophobic in nature and the HPMC-C_12_ polymers possess hydrophobic moieties, both of which act as sites for robust non-covalent, hydrophobic interactions between the drug molecules and the hydrogel components. As the standard clinical dosing of semaglutide for type 2 diabetes is ∼1 mg weekly, we anticipated that a single administration capable of providing upwards of four months of continuous therapy must contain ∼20 mg of semaglutide in a volume relevant for subcutaneous administration (e.g., generally 0.5–2 mL). We have previously reported that these injectable, biocompatible PNP hydrogel materials are capable of facile loading of diverse protein cargo at relevant concentrations with high efficiency, and enable tunable cargo delivery timeframes (*33, 37, 40–44*).

PNP hydrogels are fabricated by mixing a solution of dodecyl-modified hydroxypropyl methylcellulose (HPMC-C_12_) with a solution of biodegradable nanoparticles composed of poly(ethylene glycol)-*b*-poly(lactic acid) (PEG-PLA NPs) comprising the pharmaceutical agent of interest (Figure 1) (*37*). To form these hydrogels, a solution of PEG-PLA NPs is loaded into one sterile syringe, while a solution of HPMC-C_12_ is loaded into a separate syringe and the two solutions are mixed with a sterile Luer lock elbow mixer (Figure 2A). Upon mixing, the strong yet dynamic PNP interactions between the HPMC-C_12_ polymers and the PEG-PLA NPs constitute physical cross-linking that drives formation of a solid-like hydrogel material with 100% loading efficiency of the pharmaceutical agent, which was confirmed by dissolving fully-formulated drug-loaded hydrogels and quantifying the drug content (Figure 2A). The properties of these PNP hydrogels can be easily tuned through alteration of the formulation (i.e., the concentration of HPMC-C_12_ and PEG-PLA NPs), enabling facile modulation of the rates of hydrogel erosion and commensurate payload release. We have also previously reported that these hydrogels are exceptionally well tolerated in numerous indications and do not exhibit swelling when exposed to physiologic conditions (*34, 48*). These PNP hydrogels, therefore, can be readily optimized to target desirable release characteristics for pharmaceutical agents of interest in a broad range of therapeutic applications.

**Fig. 2.**
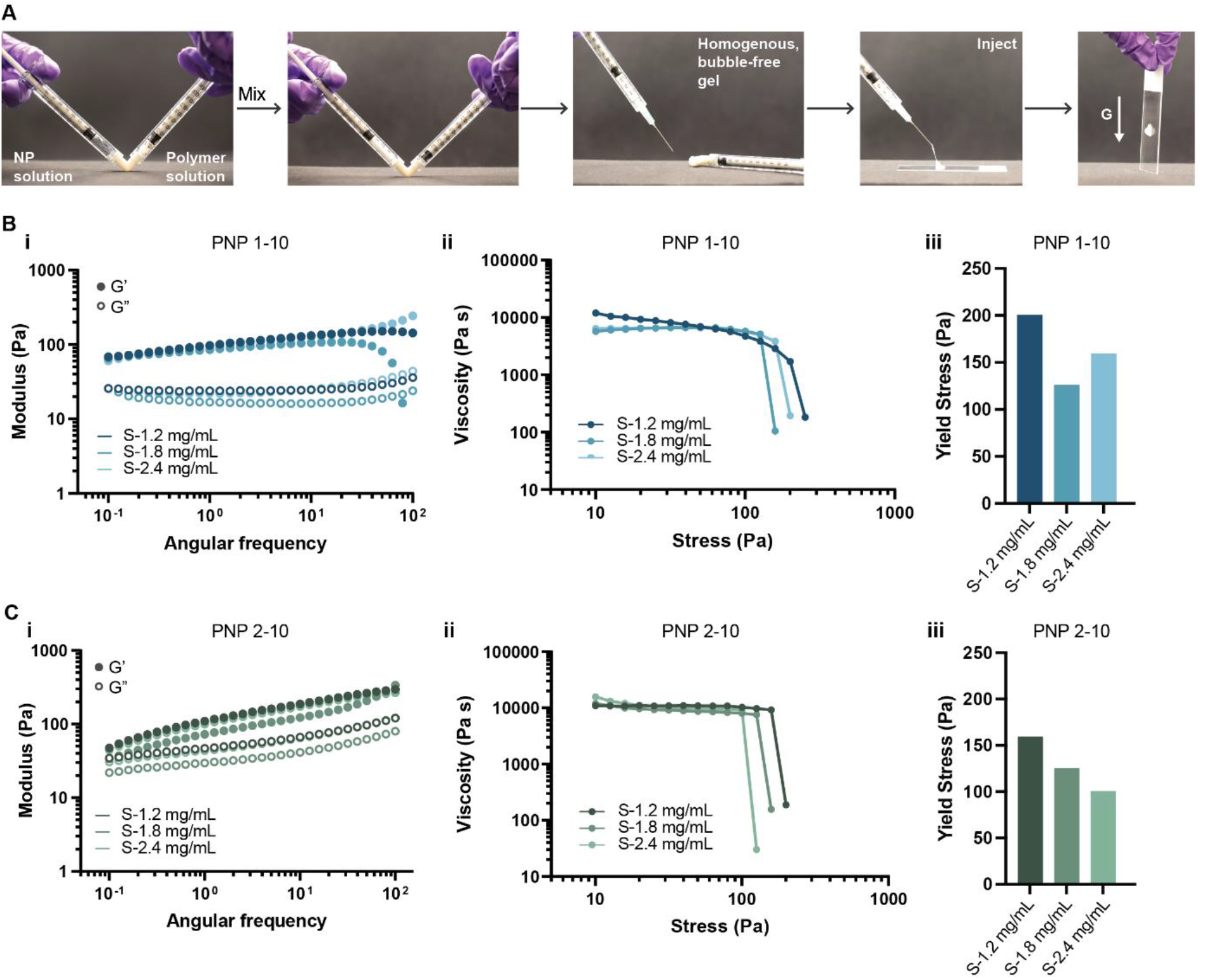
Preparation and characterization of GLP-1 RA loaded PNP hydrogel formulations. A) PNP hydrogels are prepared by mixing a solution of hydrophobically-modified HPMC (right syringe) with a solution of PEG-PLA nanoparticles and GLP-1 RAs (left syringe) using a Luer lock mixer. After mixing, a homogenous, bubble-free, solid-like PNP hydrogel is formed (right syringe). Owing to their dynamic crosslinking, PNP hydrogels are injectable through clinically relevant, high gauge needles and rapidly self-heal after injection. B) Rheological characterization of PNP-1-10 hydrogel formulations: (i) frequency-dependent oscillatory shear sweep, (ii) stress-dependent oscillatory shear sweep, and (iii) yield stress obtained from stress sweep. C) Rheological characterization of PNP-2-10 hydrogel formulations: (i) frequency-dependent oscillatory shear sweep, (ii) stress-dependent oscillatory shear sweep, and (iii) yield stress obtained from stress sweep.

### Rheological characterization of GLP-1 RA loaded PNP hydrogels

We prepared two PNP hydrogel formulations: PNP-1-10 and PNP-2-10, where the first number denotes the weight % (wt%) of polymer and the second number denotes the wt% of nanoparticles (n.b., the remaining mass is buffer). For all in vitro studies, PNP hydrogel formulations comprising semaglutide were denoted with an “S” followed by their corresponding drug loading (e.g., S-1.8 mg/mL), while liraglutide formulations were denoted with an “L” followed by their corresponding drug loading (e.g., L-1.8 mg/mL). For all in-vivo results, PNP hydrogel formulations comprising semaglutide were denoted with an “S” followed by their corresponding dosage (e.g., S-0.9 mg), while liraglutide formulations were denoted with an “L” followed by their corresponding dosage (e.g., L-0.9 mg). We targeted a GLP-1-RA concentration and dosage relevant to human equivalent dosing of semaglutide (clinical dosing of semaglutide is ∼1 mg weekly) (*49, 50*), for evaluation in a rat model of insulin-impaired diabetes. In spite of the higher dose of liraglutide required for clinically relevant effect in T2D (1.8 mg/day for liraglutide compared to 1 mg/week for semaglutide), we formulated materials with equal doses of liraglutide and semaglutide to make mechanistic comparisons on incorporation and release of the peptides from the PNP hydrogels.

The rheological properties of PNP hydrogels with different loadings of both liraglutide and semaglutide were evaluated to ensure that the cargo loading did not alter the hydrogel properties integral for injectability, depot formation, and depot persistence (Figure 2B-C, Figure S2). Frequency-dependent oscillatory shear rheological testing showed that PNP hydrogel formulations exhibit solid-like behavior (G′ > G′′; tan(δ) = 0.1–0.5) across the entire range of frequencies evaluated within the linear viscoelastic regime of these materials, meaning that all PNP hydrogel formulations demonstrate mechanical properties essential for depot formation (Figure 2Bi, Figure 2Ci). Moreover, the stiffness (G′) of these materials increased with higher polymer content. Stress-controlled oscillatory shear measurements were used to evaluate yield stress and flow behavior of these materials (Figure 2Bii/iii and Figure 2Cii/iii). Only slight differences in yield stress values were observed between 1-10 and 2-10 formulations, whereby yield stress increased with increasing polymer content and negligibly lowered with the incorporation of either liraglutide or semaglutide drug molecules. At stresses below their yield stress, these materials do not flow; however, when the stress exceeds their yield stress, the materials flow and the observed viscosity drops dramatically (Figure 2B and C). Additionally, all PNP hydrogel formulations displayed shear-thinning behavior without fracture with increasing shear rates. The high degree of shear-thinning and rapid self-healing behavior observed for these PNP hydrogels loaded with GLP-1 RAs not only enables injection through clinically-relevant, small-diameter needles (*45*), but also prevents significant burst release immediately after injection on account of rapid depot formation in the body (*33, 43, 44*). Furthermore, the robust yield stress behavior exhibited by these materials (*τ*_*y*_>100 Pa) is crucial for formation of persistent depots in the subcutaneous space upon administration (*48*).

### In vitro release kinetics of GLP-1 RAs from PNP hydrogel formulations

Next, we investigated release behaviors of semaglutide and liraglutide from PNP hydrogel formulations in vitro. To study these behaviors, drug-loaded hydrogels were loaded into capillaries by injection and PBS was added to mimic infinite sink conditions at physiological temperature. Our capillary system was designed to minimize the surface-area-to-volume ratio of the hydrogels to reduce erosion effects from exposure to the infinite sink conditions and potential hydrogel disruption during sample collection. Aliquots of the surrounding medium were taken and the concentration of either semaglutide or liraglutide was quantified by ELISA to measure the release kinetics of each drug from the hydrogel formulations over time (Figure 3A). At the end of the study, the remaining hydrogel was dissolved and the retained cargo quantified by ELISA.

**Fig. 3.**
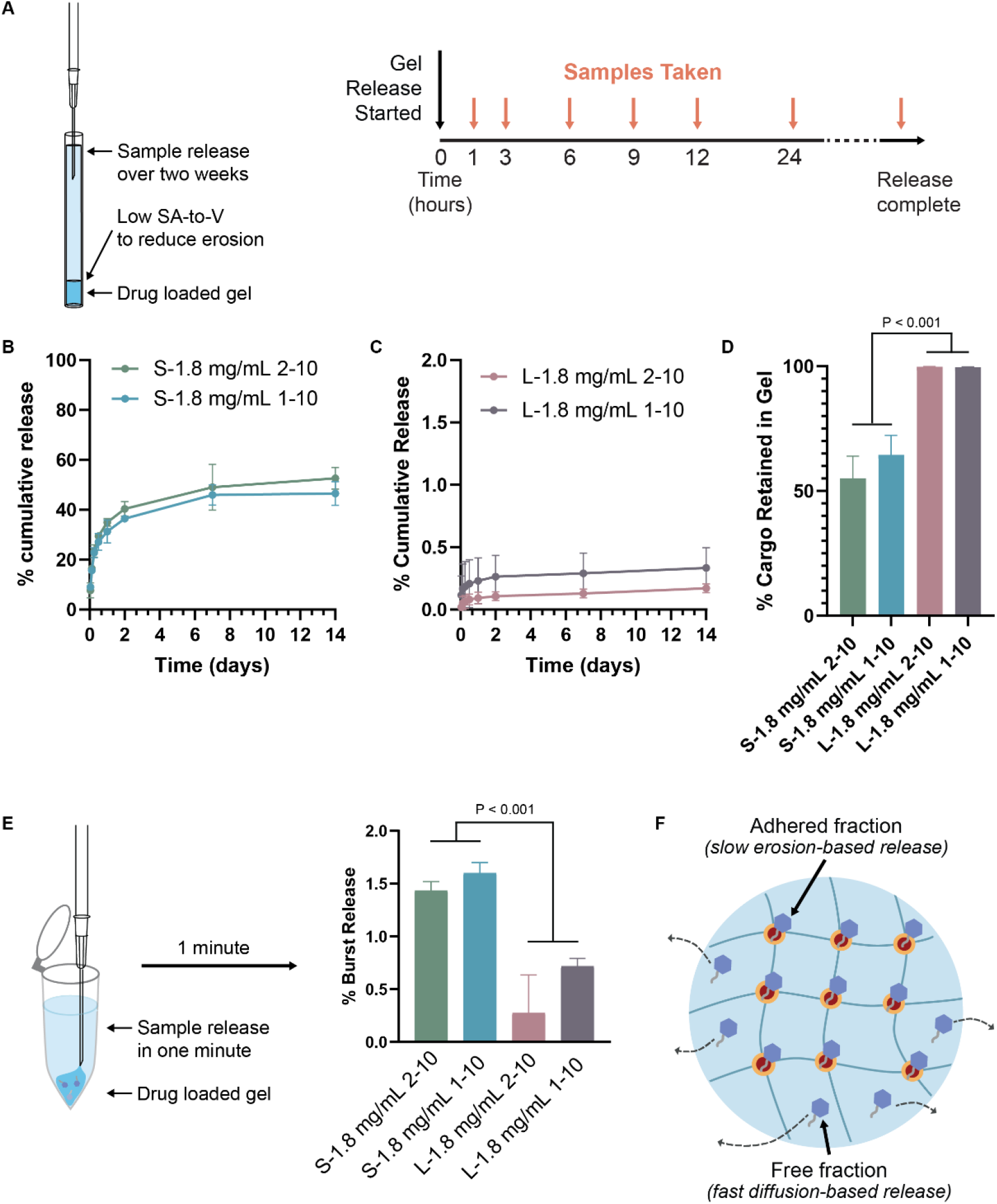
In vitro evaluation of PNP hydrogel formulation release kinetics. A) Schematic of the in vitro release assay of GLP-1 RAs from PNP hydrogels immersed in saline over two weeks. In vitro release assays were designed to minimize hydrogel erosion by minimizing the surface area-to-volume of the hydrogel. B) In vitro release profiles showing the % cumulative release of semaglutide from 1-10 and 2-10 hydrogel formulations, over the course of two weeks. C) In vitro release profiles showing the % cumulative release of liraglutide from 1-10 and 2-10 hydrogel formulations, over the course of two weeks. D) %Mass of cargo retained in the PNP hydrogel formulations during the release assay after the two-week release assay. E) Schematic of the in vitro burst release assay of the GLP-1 RAs from PNP hydrogel formulations. Samples were directly injected into saline and the supernatant saline was collected after one minute. Samples were analyzed using either semaglutide or liraglutide specific ELISA assays. The bar graph shows the % burst release for each of the hydrogel formulations after one minute. F) Schematic illustrating the different release mechanisms of cargo from the PNP hydrogels.

We compared the release of semaglutide (S-1.8 mg/mL) and liraglutide (L-1.8 mg/mL) from both PNP-1-10 and PNP-2-10 hydrogel formulations over the course of two weeks. For semaglutide in either hydrogel formulation, a clear plateau in cargo release was reached after the first week of the study, whereas liraglutide exhibited negligible release throughout the study indicative of a zero-order release profile. For semaglutide formulations, the PNP-1-10 formulation released 50 ± 5% of the entrapped cargo over the two-week study, while PNP-2-10 formulation released 54 ± 6% of cargo over the same timeframe (Figure 3B). In contrast, PNP-1-10 and PNP-2-10 hydrogel formulations comprising liraglutide released only 0.4 ± 0.2 % and 0.2 ± 0.04 % of their cargo, respectively, over the course of the two-week study (Figure 3C). From these studies, we observed that only approximately 50% of the semaglutide was retained in these hydrogels, while nearly 100% of the liraglutide was retained over prolonged timeframes in this release format where hydrogel erosion is negligible over time (Figure 3D; *P*<0.001 for comparison of semaglutide and liraglutide release).

Gastrointestinal adverse effects are common with GLP-1 RA therapy and are related to abrupt increases in drug plasma levels, which is clinically mitigated by gradual up-titration. Thus, we sought to evaluate burst release of these cargo from the PNP hydrogels, which may lead to undesirable acute high drug exposure. We established an assay in which we injected drug-loaded hydrogels directly into PBS to recapitulate the process of transcutaneous injection into the subcutaneous space, and investigated the level of burst release within the first minute after injection. The amount of released drug was quantified by ELISA one minute after injection (Figure 3E). In this assay, the S-1.8 mg/mL 2-10 formulation released 1.4 ± 0.09 % of cargo, while the S-1.8 mg/mL 1-10 formulation released 1.6 ± 0.08 % of cargo. In contrast, liraglutide formulations displayed significantly less burst release, with the L-1.8 mg/mL 2-10 formulation releasing 0.3 ± 0.4 % of cargo and the L-1.8 mg/mL 1-10 formulation releasing 0.7 ± 0.1 % of cargo. Similar to the in vitro release studies (Figure 3B-D), the semaglutide-loaded hydrogels showed a significantly greater initial burst release compared to liraglutide-loaded hydrogels (*P*<0.001 for comparision of semaglutide and liraglutide release).

As we see a higher amount of burst release and lower proportion of drug retained over prolonged timeframes for semaglutide, we hypothesized that there is likely a substantial fraction of “free” cargo (i.e., not adhered to the PNP hydrogel matrix) that undergoes fast, diffusion-based release. Such “free” drug would be expected to be subject to diffusional release over time, whereas “bound” drug would only be released by hydrogel erosion (*51*), which is severely limited in this capillary release model (Figure 3F). In contrast, liraglutide exhibited significantly higher retention within the PNP hydrogels than semaglutide (*P* = 0.0003; L-1.8 mg/mL 1-10 vs. S-1.8 mg/mL 1-10 formulations). As liraglutide formulations exhibited almost complete cargo retention and negligible burst release, we hypothesize that essentially all of the cargo is adhered to the PNP hydrogel structure and will undergo slow erosion-based release (Figure 3F). While liraglutide contains a C_12_ linker with a single carboxylic acid, semaglutide contains a C_18_ linker with two carboxylic acid moieties, making it significantly more hydrophilic (*23*). These physicochemical differences may explain the observed differences in interactions and resulting release behavior of the drugs with the hydrophobic interface of the PEG-PLA NPs of the PNP hydrogels.

### In-vivo pharmacokinetics of hydrogel-based GLP-1 RA formulations

Prolonged delivery of GLP-1 RAs affords the opportunity for a months-long-acting T2D therapy that provides steady levels of the incretin hormone to reduce the burden of maintaining glucose homeostasis. To assess the performance of our hydrogel-based formulations in vivo, we conducted pharmacokinetic and pharmacodynamic studies in a rat model of insulin-impaired diabetes (Figure 4). Diabetes was induced using a combination of streptozotocin (STZ) and nicotinamide (NA) in rats, which is a mimic of a T2D-like phenotype (*52, 53*).

**Fig. 4.**
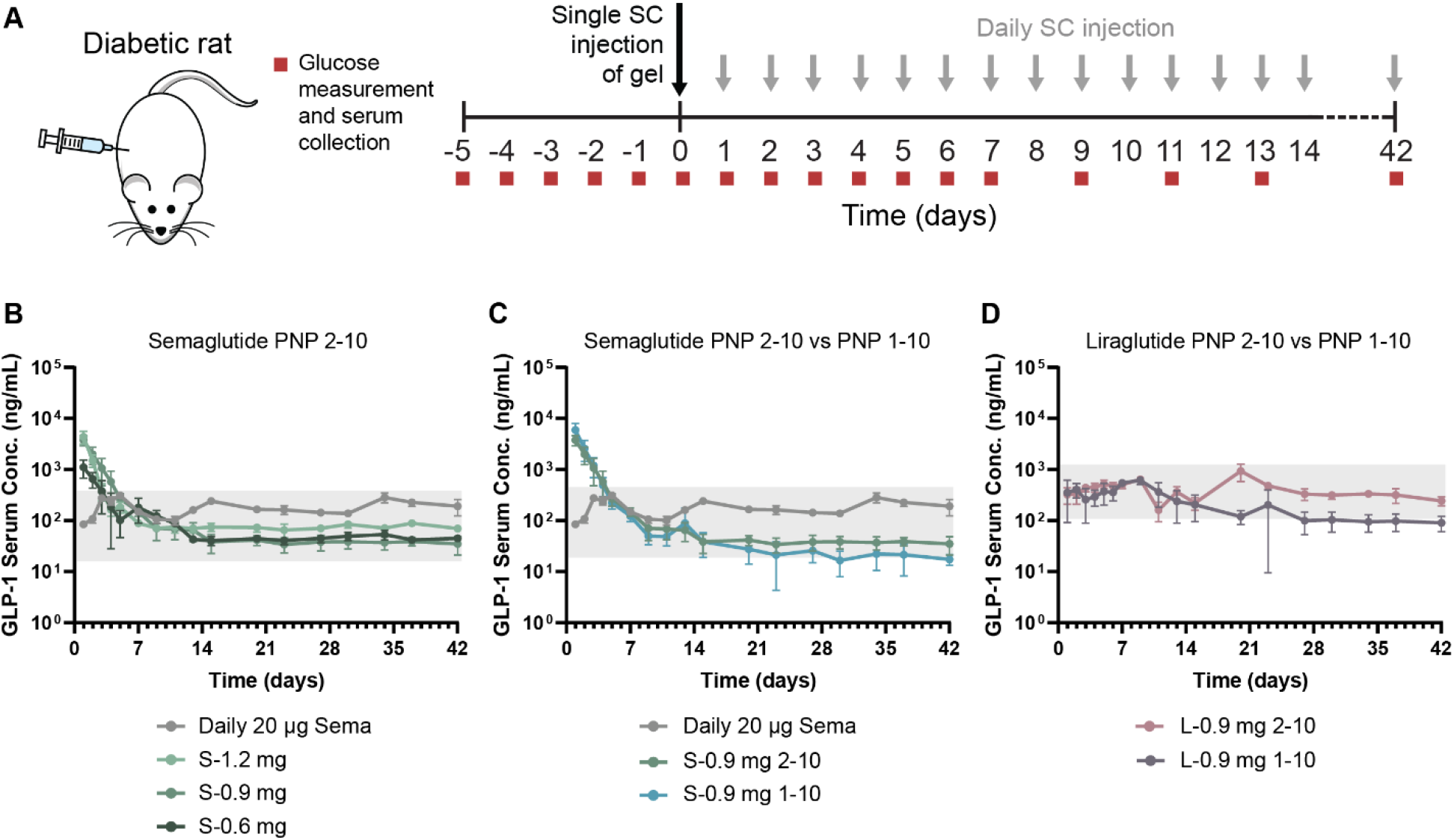
In-vivo evaluation of PNP hydrogel formulation release kinetics in diabetic rats. Single dose, GLP-1 RA hydrogel treatment prolongs the delivery of semaglutide and liraglutide for six weeks, compared to a daily 20 µg semaglutide injection. A) Treatment schedule and timing of blood glucose measurements and serum collection for analysis. Diabetic rats received either a single SC injection of hydrogel, or daily SC bolus injection of either PBS or 20 µg daily injections of semaglutide. B) Pharmacokinetics in diabetic rats. Fasted male diabetic rats (n = 5–6 received subcutaneous administration of i) high loading (S-1.2 mg) of semaglutide, ii) medium loading (S-0.9 mg) of semaglutide and iii) low loading (S-0.6 mg) of semaglutide in 2-10 formulations. C) Comparison of 2-10 versus 1-10 hydrogel formulation pharmacokinetics, with the same semaglutide loading (S-0.9 mg). D) 2-10 versus 1-10 liraglutide hydrogels at a loading of 0.9 mg.

Following the onset of diabetes according to literature procedures (*53, 54*), rats were fasted overnight to ensure an average starting blood glucose (BG) level of approximately 130 mg/dL to 200 mg/dL (*55*). Fasting BG levels were then measured once a day for five days before treatment and rats were considered diabetic when at least three out of five fasting BG measurements, taken over the course of five days, fell between 130–200 mg/dL. Animals with BGs lower than 130 mg/dL or above 200 mg/dL, were excluded from the study (Figure 4A). Initially, we validated previously reported pharmacokinetic parameters (*23*) in diabetic rats (i.e., elimination half-life, volume of distribution, bioavailability, and absorption rate) by conducting a 24-hour pharmacokinetic (PK) study following intravenous (IV) administration of a 20 µg dose of semaglutide in a saline vehicle (Figure S3). After validating the pharmacokinetics, we conducted an IV glucose tolerance test (IV GTT) to ensure similar glucose response when grouping the rats into treatment groups. Based on the IV GTT, rats with similar glucose tolerance profile were paired, then these rats were randomized into treatment groups (Figure S4). We then compared the PK following subcutaneous administration of four treatment regimens: (i) repeated daily 20 µg injections of semaglutide in a saline vehicle to mimic current clinical practice (*23*), (ii) single semaglutide hydrogel formulations, (iii) single liraglutide hydrogel formulations, and (iv) repeated daily PBS injections as an untreated control (n = 5–6 for each treatment group). Semaglutide has a half-life of 7 days in humans at a typical dose of 0.5–1 mg (*56*), and PK modeling shows that daily 20 µg dosing in rats recapitulates the current clinical treatment regimen for patients (*32*). In contrast, the single hydrogel treatment groups represent our approach to long-acting GLP-1 RA delivery. For the first seven days after the start of treatment, and three times a week thereafter, serum was collected for PK analysis and BG was measured (Figure 4A).

The serum concentrations of either semaglutide or liraglutide were measured over time following the subcutaneous administration of each treatment by ELISA to assess the pharmacokinetic profiles of each hydrogel-based formulation. We hypothesized that our hydrogel formulations would maintain therapeutically relevant concentrations of semaglutide and liraglutide, comparable to daily 20 µg semaglutide administration, for six weeks in this rat model. Indeed, all six hydrogel formulations we evaluated in this study effectively maintained relevant concentrations of semaglutide or liraglutide throughout the duration of the six-week-long study (Figure 4; n.b., the window equivalent to human relevant serum concentrations for each drug is indicated by the grey box).

One important metric for adverse effects of GLP-1 RA therapies (primarily gastrointestinal discomfort that is most prevalent when initiating treatment) is C_max_, which is the maximum concentration of GLP-1 RA in the serum following treatment. In our studies, daily 20 µg semaglutide administration yielded a C_max_ of 280 ± 60 ng/mL (observed at day 34), though serum levels were relatively stable over the course of the treatment period with no clear peaks in the PK profile (*56*). In contrast, semaglutide-loaded hydrogels reached very high C_max_ values within one day of treatment, followed by a continuous decrease in serum concentrations over the course of the first week to a steady-state that remained throughout the duration of the study. The C_max_ values for each treatment group 1110 ± 430 ng/mL (S-0.6 mg 2-10), 3800 ± 900 ng/mL (S-0.9 mg 2-10), and 4400 ± 1200 ng/mL (S-1.2 mg 2-10) (Figure 4B, Table S2, S3). There was a trend towards higher observed C_max_ values with increasing dose of semaglutide in these hydrogel formulations, and the highest dose S-1.2 mg 2-10 formulation exhibited significantly higher C_max_ values than the S-0.6 mg 2-10 formulation (*P* = 0.0003; Table S2 and S3). Similarly, the S-0.9 mg 1-10 formulation exhibited a higher C_max_ (5900 ± 2100 ng/mL) compared to the corresponding S-0.9 mg 2-10 formulation (3800 ± 900 ng/mL, *P* = 0.09; Table S2 and S3). It is important to note that while the lowest dose hydrogel formulation (S-0.6 mg 2-10) exhibited the lowest C_max_ compared to the other hydrogel formulations (1100 ± 430 ng/mL), this value was much higher than the C_max_ of the daily 20 µg semaglutide treatment group (280 ± 60 ng/mL, *P* = 0.003; Table S2 and S3). While the C_max_ values observed within the first day for all hydrogel-based semaglutide formulations might be expected to cause nausea and suppressed food intake in rodents, no changes to animal behavior were observed throughout the study.

Contrary to the PK behaviors observed for the semaglutide-based formulations, both liraglutide-based hydrogel formulations quickly established steady-state serum concentrations and did not exhibit any peaks in their pharmacokinetic profiles, indicating a more tolerable profile. The L-0.9 mg 1-10 hydrogel formulation exhibited a C_max_ of 620 ± 110 ng/mL (observed on day 9), while the L-0.9 mg 2-10 hydrogel formulation exhibited a C_max_ of 930 ± 350 ng/mL (observed on day 20) with relatively steady plasma concentrations throughout the duration of the study (Figure 4D, Table S2). These observations are consistent with the release and retention data described above, whereby the large fraction of “free” semaglutide drug observed within these hydrogel-based formulations would be expected to be released rapidly, thereby contributing to elevated C_max_ values at early timepoints. In contrast, liraglutide was completely retained within the hydrogel depot and would be expected to only be released slowly over time with depot dissolution.

Another important metric for effective therapy is maintenance of appropriate steady-state GLP-1 RA serum concentrations. Each of the six hydrogel-based GLP-1 RA formulations achieved steady-state kinetics throughout the six-week study, with the semaglutide-loaded 1-10 and 2-10 formulations reaching steady-state within one week of treatment, while the liraglutide-loaded 1-10 and 2-10 hydrogel formulations reaching steady-state within one day of treatment (Table S2, S4). Consistent with the trends observed for C_max_, a higher semaglutide dose yielded higher C_steady-state_ values, whereby the S-1.2 mg 2-10 and S-0.6 mg 2-10 formulations achieved steady-state serum concentrations of 75 ± 10 ng/mL and 66 ± 50 ng/mL, respectively (*P* = 0.8, Table S2 and S4).

The hydrogel formulation (i.e., 1-10 vs 2-10 hydrogels) had less of an impact on steady-state serum concentrations for both semaglutide and liraglutide. Semaglutide-based S-0.9 mg 1-10 and S-0.9 mg 2-10 hydrogel formulations reached similar C_steady-state_ values of 40 ± 40 ng/mL and 60 ± 40 ng/mL, respectively (*P* = 0.3, Table S2 and S4). Similarly, liraglutide-based L-0.9 mg 1-10 and L-0.9 mg 2-10 hydrogel formulations reached C_steady-state_ values of 270 ± 190 ng/mL and 410 ± 200 ng/mL, respectively (*P* = 0.02, Table S2 and S4).

### In-vivo pharmacodynamics of hydrogel-based GLP-1 RA formulations

Next, we examined the ability of the hydrogel-based long-acting GLP-1 RA formulations to reduce average BG after each six-week treatment regimen. In this study, all GLP-1 RA treatments resulted in a significant reduction in average BG over the course of the study, whereas untreated controls experienced no significant change in average BG (Table S5). While BG dropped more noticeably during the first two weeks of the study for the semaglutide hydrogel treatment groups, the reduction was maintained throughout the six-weeks. The daily dosing of 20 µg semaglutide resulted in a 14 ± 4% reduction in average BG over the course of the six-week study. While a single administration of the S-0.6 mg 2-10 (15 ± 7% reduction) and S-0.9 mg 2-10 (18 ± 10% reduction) hydrogel treatments were just as effective at lowering the average BG levels as the daily 20 µg semaglutide administrations (*P* = 0.758 and *P* = 0.586, respectively, Figure 5A), a single higher dose S-1.2 mg 2-10 (27 ± 2% reduction) hydrogel treatments was significantly more efficacious than the daily 20 µg semaglutide dose (*P* = 0.0002, Figure 5A). Moreover, the S-0.9 mg 1-10 hydrogel formulation resulted in a mean reduction in average BG of 29 ± 4%, similar to the S-0.9 mg 2-10 hydrogel formulation, and was also significantly more efficacious than the daily 20 µg semaglutide treatment (*P* = 0.0043, Figure 5A). The liraglutide-based hydrogel formulations reduced BG comparable to daily semaglutide administration, with the L-0.9 mg 1-10 and L-0.9 mg 2-10 treatments resultin in mean reductions in average BG of 20 ± 6% (*P* = 0.111) and 22 ± 8% (*P* = 0.0929), respectively.

**Fig. 5.**
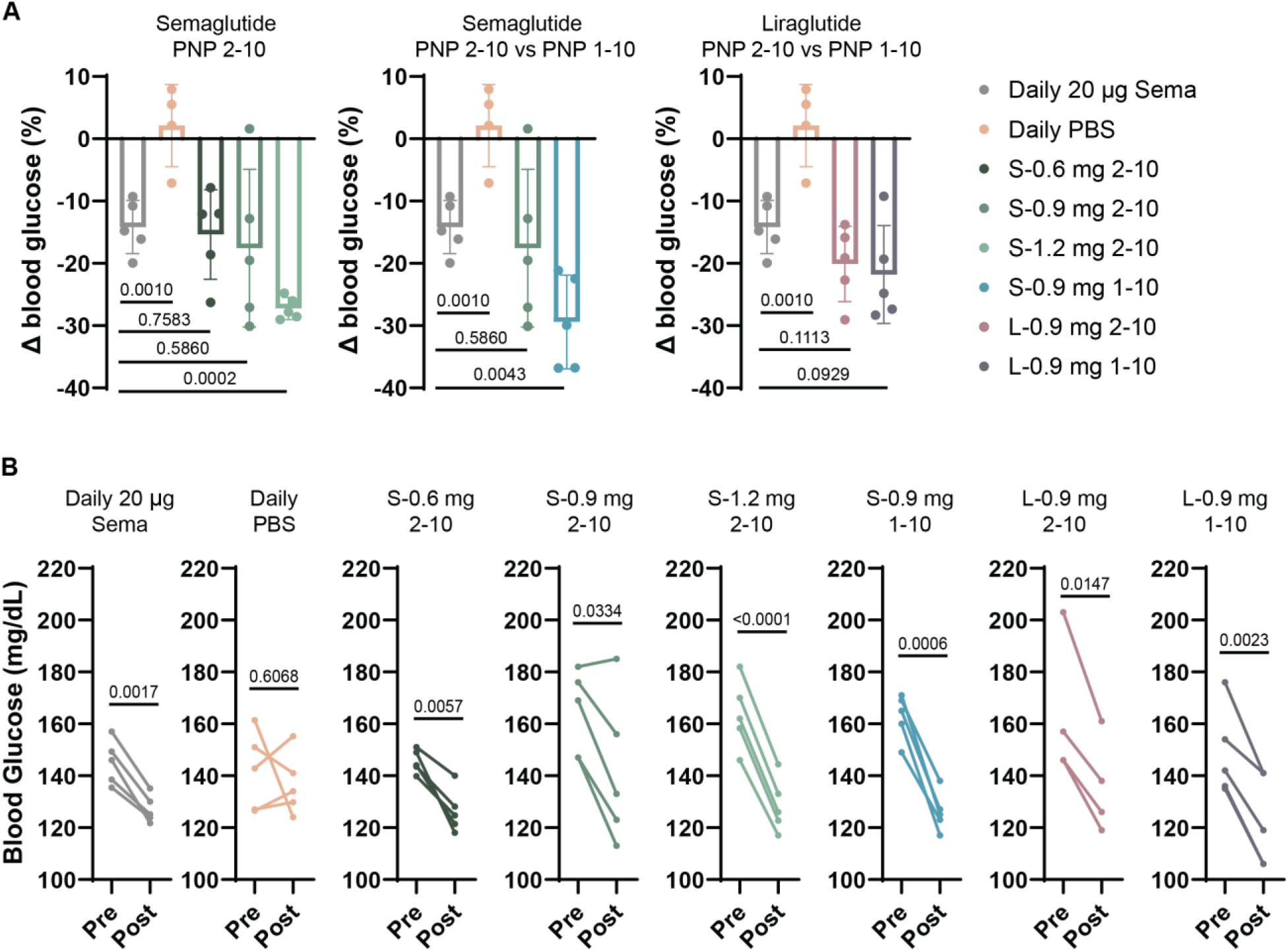
Effect of PNP hydrogel formulations on blood glucose in diabetic rats. A single administration of GLP-1 RA hydrogels reduces the BG of type 2-like diabetic male rats over the course of six weeks and are similar or more effective than a treatment regimen consisting of daily semaglutide bolus injections. A) Change in BG over 6 weeks following each treatment group regimen (n = 5–6). B) Individual pre- and post-treatment regimen blood glucose values for individual rats in each treatment group (n = 5–6). *P* values were determined using a one-way ANOVA with Dunnett’s multiple comparisons test and using an unpaired, two-tailed t-test. Table S5 lists the *P* values of each treatment group compared with the PBS control.

We also compared the effect of our hydrogel-based GLP-1 RA formulations with daily semaglutide treatment on body weight throughout the study (Figure 6). These young growing rats, which increased in body weight over the course of 42-days, exhibited a less pronounced weight gain and a resulting lower body weight at the end of the study when receiving daily semaglutide (41 ± 10% weight gain) compared to a saline control (65 ± 13% weight gain) (*P* = 0.0114, Table S6). Similar to our observations above regarding blood glucose, single hydrogel-based treatments resulted in similar weight management as daily semaglutide (Figure 6A). For the three semaglutide-based 2-10 hydrogel formulation treatment groups, we observed that the lowest dose S-0.6 mg 2-10 treatments exhibited the greatest increase in weight (50 ± 11%; *P* = 0.2066 w.r.t. daily semaglutide), while the higher dose S-0.9 mg 2-10 and S-1.2 mg 2-10 treatments exhibited average increases in weight of 38 ± 8% and 42 ± 6% that were similar to the daily semaglutide treatment (*P* = 0.6466 and *P* = 0.8866, w.r.t. daily semaglutide). While the S-0.9 mg 2-10 formulation (38 ± 8%; *P* = 0.6466 w.r.t. daily semaglutide) and the L-0.9 mg 1-10 formulation (40 ± 16%; *P* = 0.8945 w.r.t. daily semaglutide) resulted in the best overall weight management, comparable to daily semaglutide treatment, all hydrogel-based formulations were highly effective at mitigating weight gain in these young, lean growing rats (Figure 6).

**Fig. 6.**
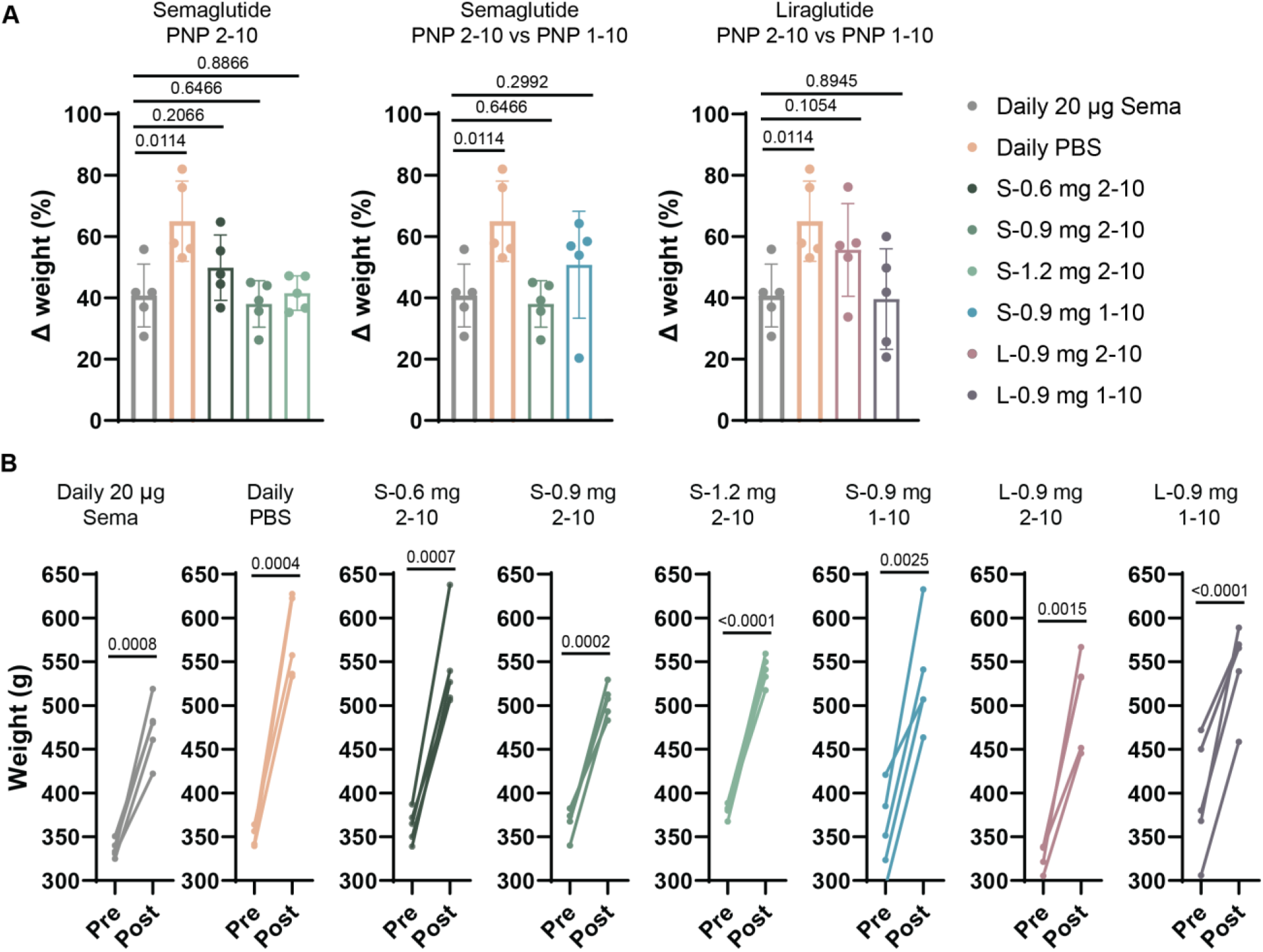
Effect of PNP hydrogel formulations on weight in diabetic rats. A single administration of GLP-1 RA hydrogels reduces the overall weight gain in type 2-like diabetic male rats over the course of six weeks. A) Change in weight over 6 weeks of each treatment group (n = 5–6). B) Individual pre- and post-treatment weight for individual rats in each treatment group (n = 5–6). *P* values were determined using a one-way ANOVA with Dunnett’s multiple comparisons test and using an unpaired, two-tailed t-test. Table S5 lists the *P* values of each treatment group compared with the PBS control.

### Biocompatability of hydrogel-based GLP-1 RA therapeutics

Since these hydrogel-based long-acting GLP-1 RA therapeutics are novel drug product candidates, we also sought to assess their biocompatibility. The PNP hydrogels themselves have been shown previously to be highly biocompatible and non-immunogenic in mice, rats, pigs and sheep in various therapeutic contexts (*33–39, 41, 44*). Here, we sought to conduct histopathological analysis at the end-point of the rat studies outlined above. A blinded assessment of the histopathology of the liver and kidney of treated animals was conducted 6 weeks after treatment (Figure S5 and S6). The hydrogel-based treatments were shown to be well tolerated, exhibiting no observable differences in liver and kidney compared to both untreated animals and animals receiving daily semaglutide treatment. These results are promising as they indicate that the hydrogel-based treatments are highly biocompatible.

### Modeling of long-acting liraglutide-hydrogel pharmacokinetics in humans

Based on all of the data described above, the L-0.9 mg 1-10 and 2-10 hydrogel formulations exhibited the most consistent and favorable pharmacokinetics, reaching steady state after just one day and maintaining stable and therapeutically relevant concentrations for the duration of the 42-day study. Yet, beyond evaluating the pharmacokinetics (PK) and pharmacodynamics (PD) of hydrogel-based GLP-1 RA therapies, simple pharmacokinetic modeling can be used to predict drug release kinetics in more clinically relevant large animals and in humans, while also aiding in the design of biomaterial depot technologies (*42*). Modeling can inform the optimization of the PNP hydrogel platform for delivery of cargo with similar physicochemical characteristics but different pharmacokinetic characteristics. We have previously reported a simple PK model for depot-based biopharmaceutical formulations following SC administration, including in PNP hydrogels, based on a two-compartment model that takes into account species-relevant dosing and physiological values for GLP-1 RA PK (Figure 7A) (*57*). Here, we used this approach to model the PK for liraglutide in rats following prolonged release from the PNP hydrogel depot and leverage the observed in-vivo release behaviors to predict the PK of these hydrogel-based liraglutide treatments in humans.

**Fig. 7.**
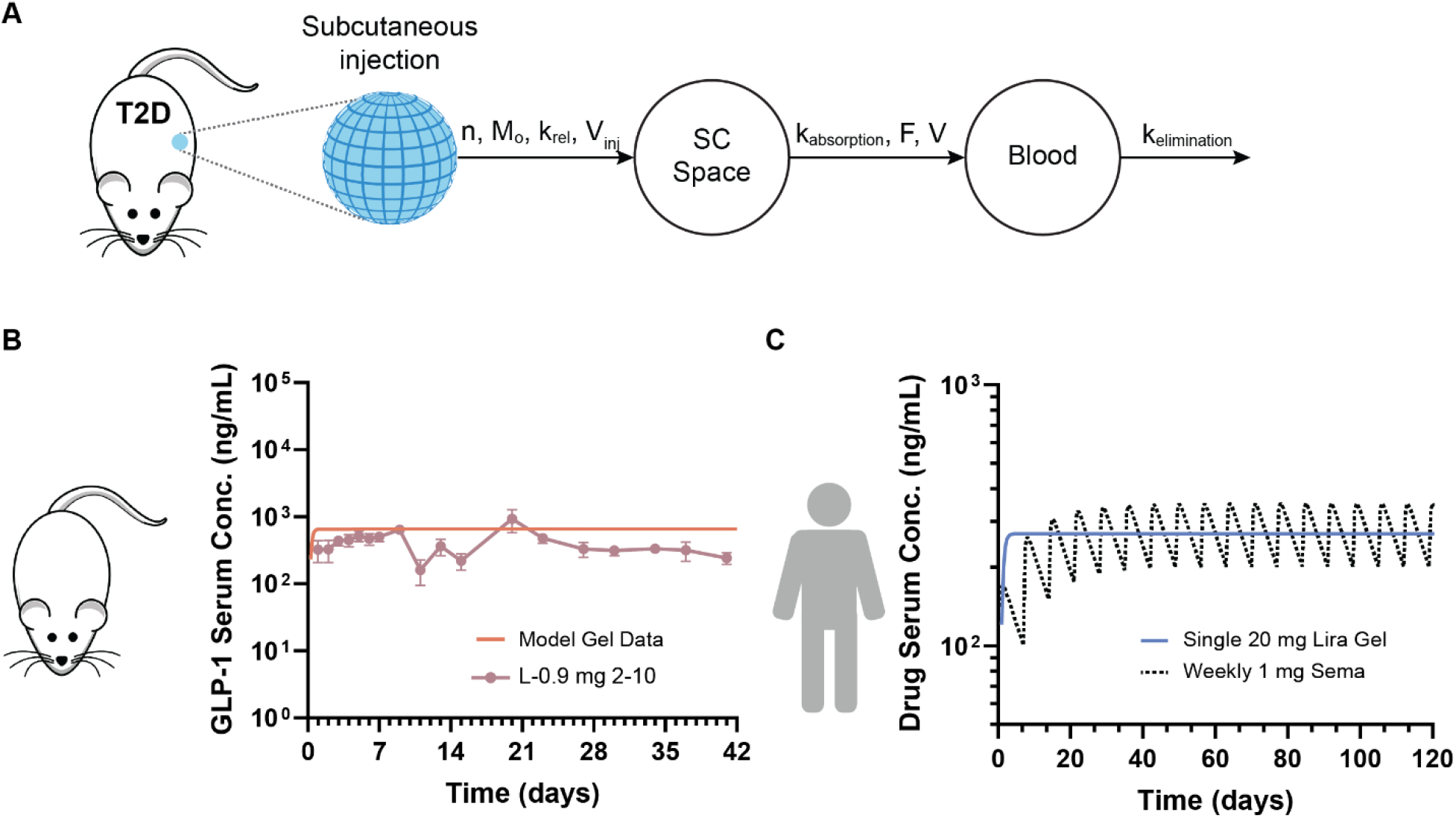
Pharmacokinetic modeling of hydrogel-based liraglutide formulations in humans. A) Scheme showing a two-compartment pharmacokinetic model of drug release from a subcutaneous hydrogel depot. In this model, k_rel_ and n are the release parameters of the molecular cargo from the hydrogel depot. B) Pharmacokinetics of 0.9 mg liraglutide 1-10 hydrogels in type 2-like diabetic rats following SC administration indicate a single 500 µL injection of PNP-1-10 hydrogels can maintain therapeutic concentrations of liraglutide for 6 weeks when typical bolus treatment with semaglutide must be administered daily. PK modeling can be used to determine *in-vivo* release characteristics for our hydrogel (k_rel_) with known parameters for SC absorption and elimination of liraglutide in rats. C) Hydrogel release characteristics *in-vivo* can be used to estimate PK in humans by using known PK parameters for absorption and elimination and typical drug dosing. Human PK modeling suggests that a single 1.25 mL injection of liraglutide-loaded PNP-1-10 hydrogels can potentially maintain therapeutically relevant concentrations of liraglutide for up to four months, matching serum concentrations achieved through weekly injections of commercial semaglutide formulations according to standard dosing regimens.

The liraglutide PK for the L-0.9 mg 1-10 was modeled using the half-life of absorption from the SC space (t_1/2, SC_ ∼ 0.16 h) and the known half-life of elimination from systemic circulation in rats (t_1/2, serum_ ∼0.167 days) (Table S1) (*58*). This simple two compartment model sufficiently recapitulates the observed liraglutide PK following release from the SC hydrogel depot, suggesting that the model adequately estimates the mass transport processes occurring in vivo. This modeling thereby provides insight into the species-independent release characteristics of these materials that enable scaling of the PK to human dosing, assuming full chemical and physical stability of the drug molecule and complete release from the hydrogel depot. For the 500 µL injection volume administered in the rats, we observe a rate of liraglutide release of ∼2% per day, which is comparable the erosion rates observed for 100 µL administrations of PNP-1-10 hydrogels observed previously in mice (*48*). These erosion rates correspond to a liraglutide release rate of ∼0.8% per day from a clinically-relevant 1.25 mL subcutaneous injection volume administered in humans. Indeed, the 2.5-fold larger injection volume relevant for human administration will result in a 2.5-fold reduction in the release rate observed in rats (Table S7, S8). Using the expected liraglutide release rate in humans following clinically-relevant administration (0.8% per day; 1.25 mL injection volume), relevant human dosing (20 mg), and known human PK parameters for liraglutide (t_1/2, SC_ ∼ 0.25 days, t_1/2, serum_ ∼0.5417 days) (*59*), we then modeled the predicted liraglutide PK in humans following hydrogel-based treatments (Figure 7C). This modeling suggests that the 1-10 PNP hydrogel formulation can potentially maintain therapeutically relevant levels of liraglutide in systemic circulation for upwards of 120 days from a single administration. From these modeling studies, we observe that a single hydrogel-based treatment has the potential to maintain drug serum concentrations comparable to weekly administrations of semaglutide, one of the leading commercial GLP-1 RA drugs (Figure 7C).

## DISCUSSION

Patient adherence to antihyperglycemic treatment, medications is surprisingly low, described for GLP-1 RA to fall between 29% and 54%, resulting in suboptimal T2D management carrying increased risk of stroke, heart and kidney disease, amputations and blindness (*60*). For drugs with short half-lives, poor compliance with prescribed treatment regimens reduces plasma concentrations to unsuitable levels, and multiple doses are often required to return to therapeutic plasma concentrations. Complex and/or frequent administration of treatment is one hurdle to adherence. Aligning a GLP-1 RA dosing regimen with the typical schedule of clinical visits has the potential to greatly improve compliance while enabling adjustments to a treatment plan at these visits. Indeed, a once-every-four-month therapy is likely to be ideal for clinical diabetes management, as this would align with the typical cadence with which patients visit their endocrinologist or personal care provider. Importantly, these therapies would not only treat patients with diabetes, but more broadly impact children and adults struggling with obesity.

Recently, a combination drug-device product, ITCA 650, which consists of a matchstick-sized osmotic mini-pump that is implanted surgically into subcutaneous tissue, has been shown to enable a continuous infusion of exenatide for 3-6 months (*61, 62*). While the implantation of these devices can be performed by trained health care professionals in a brief office procedure, they must be replaced or refilled for each new dosing regimen. While the long duration of exenatide therapy from these devices is promising, the invasiveness of these interventions is unfavorable for patients and the treatment is cost-prohibitive for many patients (*63*).

To address the challenge of prolonging GLP-1 RA therapy in an easy-to-implement manner, we sought to develop long-acting GLP-1 RA formulations providing months of continuous therapy from a single administration by leveraging an injectable supramolecular hydrogel capable of prolonged delivery of GLP-1 RAs. We hypothesized that one particular class of hydrogel materials, PNP hydrogels, would be particularly well suited for this purpose and would exhibit several important features: (i) facile formulation with important GLP-1 RAs such as semaglutide and liraglutide, (ii) patient convenience in the form of straightforward injectability with standard syringes and needles, (iii) excellent tolerability by maintaining consistent slow release to circumvent undersirable gastrointestinal side-effects, and (iv) four months of continuous therapy per administration. Here, we demonstrated that PNP hydrogel materials enable simple formulation of both semaglutide and liraglutide, which is facilitated by their hydrophobic fatty acid side chains, while maintaining their rheological properties and exhibiting facile injectability. Pharmacokinetic modeling of the impact of the difference in typical subcutaneous administration volumes between rats (∼0.5 mL) and humans (∼1.25 mL), which directly influences the timeframe of hydrogel erosion and drug release, coupled with the much shorter elimination half-life of GLP-1 RAs in rats (semaglutide t_1/2, serum_ ∼ 0.29 days) than in humans (semaglutide t_1/2, serum_ ∼ 7 days), indicated that a once-every-four-month therapeutic in humans would exhibit six weeks of continuous therapy in rats.

To evaluate the efficacy of our long-acting GLP-1 RA therapeutics, we used an insulin-impaired rat model of T2D. As the half-life of semaglutide is only 0.29 days in rats (*23*), the drug must be injected daily to maintain therapeutic concentrations in systemic circulation. We observed that animals treated daily with 20 µg bolus injections of semaglutide for 42 days exhibited a decrease in average BG levels and less pronounced weight gain compared to untreated control rats receiving daily PBS injections. In contrast, we demonstrated that a single administration of either PNP hydrogel-based semaglutide or liraglutide drug product candidates maintained therapeutically relevant GLP-1 RA serum concentrations throughout the 42-day study in these T2D rats. Furthermore, a single administration of select hydrogel-based GLP-1 RA formulations, including 2-10 hydrogels comprising a 0.9 mg dose of semaglutide (S-0.9 mg 2-10) and 1-10 hydrogels comprising a 0.9 mg dose of liraglutide (L-0.9 mg 1-10), led to reduced BG levels, and improved weight management, in line with daily semaglutide injections over the course of the 42-day study.

While both semaglutide and liraglutide hydrogel-based formulations were shown to improve glucose as well as body weight control, we found that semaglutide-loaded hydrogel formulations exhibited high C_max_ values within one day of treatment and high drug exposure over the first week of therapy, which is likely to be poorly tolerated in humans. Following the high initial exposure, these semaglutide-based hydrogel treatments exhibited a steady-state plateau in serum drug concentrations throughout the duration of the study. In contrast, the liraglutide-loaded hydrogel formulations exhibited no initial peak and reached steady-state serum drug concentrations within one day of treatment that persisted throughout the 42-day study. These results suggest that a significant fraction of the semaglutide is released from the hydrogels over the first week, followed by longer-term controlled release. In contrast, all of the liraglutide was found to be entrapped within the hydrogels and released in a more controlled manner as the hydrogels erode by dissolution over time, providing consistent serum exposure throughout the treatment period. These observations were corroborated by our in vitro release studies, which indicated that approximately 50% of entrapped semaglutide is released from the hydrogels within the first week when a plateau was reached, while less than 0.3% of entrapped liraglutide is released under similar conditions. It is important to note that a slow uptitration regimen is important for minimizing the risk of gastrointestinal symptoms in patients (*64*). Based on our results, our liraglutide hydrogels provide a steady state release of GLP-1 RAs within the therapeutic window, and we don’t see any adverse gastrointestinal effects in our rodent model at these concentrations. We would anticipate that if gastrointestinal effects are observed in humans, uptitration could be conducted with currently used one-week-long GLP-1 RA drug products such as Ozempic prior to using a once-every-four-month drug product such as the ones described here.

These results suggest that the semaglutide-loaded hydrogels may contain a significant fraction of “free” cargo that is released diffusively over short timeframes from the hydrogels, while the remaining “bound” fraction is released over long timeframes as the hydrogels dissolve away. Similarly, liraglutide appears to be entirely “bound” to the hydrogel structure and only releases through hydrogel dissolution and erosion. Previous literature reports have demonstrated that the fatty acid side chain of liraglutide drives the formation of more disorganized heptameric structures at micromolar concentrations (*65*), while semaglutide has been reported to form robust dimeric species at similar concentrations (*66, 67*). Additionally, semaglutide’s side chain linker, which contains two carboxylic acid moieties, is significantly more hydrophilic than that of liraglutide’s, which only contains one carboxylic acid moiety (*23*). These physicochemical differences may result in different interactions of the drugs with the hydrophobic interface of the PEG-PLA NPs that constitute the PNP hydrogels. Our observations suggest that the liraglutide may associate more strongly with the structural motifs within the PNP hydrogels to form such a high “bound” fraction, whereas the robust semaglutide dimers formed at formulation-relevant concentrations constitute the “free” fraction of drug cargo. Importantly for this discussion, semaglutide dimers are sufficiently small (*R*_H_ < 2 nm) to be released over relatively short timeframes from the PNP hydrogels on account of its comparatively large mesh size (*ξ*∼3.5 nm) (*40, 66*), commensurate with the release behaviors we observed both in vitro and in vivo.

In addition to demonstrating the pharmacokinetics and pharmacodynamics of these hydrogel-based GLP-1 RA treatments, we have leveraged compartment modeling to demonstrate that a single administration of these drug product candidates can potentially provide upwards of four months of continuous therapy in humans. Using this approach to pharmacokinetic modeling, which appropriately captured our experimental pharmacokinetic data in rats (Figure 7B, Table S7), enabled us to characterize the release kinetics of the GLP-1 RA drugs from our hydrogel depot in vivo. As the administration volume will be much higher for humans (∼1.25 mL) than rats (∼0.5 mL), modeling shows an extension of the pharmacokinetics to maintain therapeutically relevant serum concentrations of drug for 120 days as a larger hydrogel depot will take longer to erode by dissolution (Table S8). Incorporating this type of compartment modeling during the hydrogel material design process is crucial for determining what sort of release behaviors are required to achieve a desired drug PK profile and to ensure that depot technology design is relevant for translation.

This work lays the foundation for translation of advanced hydrogel technologies for the prolonged release of therapeutic peptide-based anti-diabetic and anti-obesity treatments. We have generated months-long-acting GLP-1 RA drug product candidates forming the basis for a transformational new approach to managing diabetes and obesity. Importantly, as liraglutide is the only currently approved GLP-1 RA therapy approved for children, this work will have broad impacts on a broad range of people. GLP-1 RA-loaded PNP hydrogels constitute an efficacious treatment approach that can reduce patient burden by requiring only one injection every four months. Indeed, a single administration of these hydrogel-based treatments provided blood glucose and body weight control comparable to 42 administrations of current clinical treatments in a rodent model of T2D. While this research has the potential to impact people with T2D, as they would benefit directly from a long-acting GLP-1 drug product that can reduce treatment burden and improve disease management, numerous recent studies suggest that such a product can improve the quality of treatment for people with obesity or T1D as well (*17, 68–70*). Furthermore, beyond treatments for diabetes and obesity, this work has the potential to significantly advance the development of long-acting formulations of therapeutic peptides and proteins more broadly.

## MATERIALS AND METHODS

### Materials

HPMC (meets USP testing specifications), N,N-diisopropylethylamine (Hunig’s base), hexanes, diethyl ether, N-methyl-2-pyrrolidone (NMP), dichloromethane (DCM), lactide (LA), 1-dodecylisocynate, and diazobicylcoundecene (DBU) were purchased from Sigma-Aldrich and used as received. Monomethoxy-PEG (5 kDa) was purchased from Sigma-Aldrich and was dried under vacuum prior to use. Glassware and stir bars were oven-dried at 180°C. When specified, solvents were degassed by three cycles of freeze, pump, and thaw. Semaglutide and liraglutide of pharmaceutical quality was provided by Novo Nordisk A/S.

### Preparations of HPMC-C_12_

Dodecyl-modified (hydroxypropyl)methyl cellulose (HPMC−C_12_) was prepared according to previously reported procedures (*71*). HPMC (1.0 g) was dissolved in NMP (40 mL) by stirring at 80 °C for 1 h. Once the solution reached room temperature (RT), 1-dodecylisocynate (105 mg, 0.5 mmol) and N,N-diisopropylethylamine (catalyst, ∼3 drops) were dissolved in NMP (5.0 mL). This solution was added dropwise to the reaction mixture, which was then stirred at RT for 16 h. This solution was then precipitated from acetone, decanted, redissolved in water (∼2 wt%), and placed in a dialysis tube for dialysis for 3−4 days. The polymer was lyophilized and reconstituted to a 60 mg mL^−1^ solution with sterile PBS.

### Preparation of PEG-PLA NPs

PEG−PLA was prepared as previously reported (*71*). Monomethoxy-PEG (5 kDa; 0.25 g, 4.1 mmol) and DBU (15 µL, 0.1 mmol; 1.4 mol% relative to LA) were dissolved in anhydrous dichloromethane (1.0 mL). LA (1.0 g, 6.9 mmol) was dissolved in anhydrous DCM (3.0 mL) with mild heating. The LA solution was added rapidly to the PEG/DBU solution and was allowed to stir for 10 min. The reaction mixture was quenched and precipitated by a 1:1 hexane and ethyl ether solution. The synthesized PEG−PLA was collected and dried under vacuum. Hydrogel permeation chromatography (GPC) was used to verify that the molecular weight and dispersity of polymers meet our quality control (QC) parameters. NPs were prepared as previously reported (*71*). A 1 mL solution of PEG−PLA in DMSO (50 mg mL^−1^) was added dropwise to 10 mL of water at RT under a high stir rate (600 rpm). NPs were purified by centrifugation over a filter (molecular weight cutoff of 10 kDa; Millipore Amicon Ultra-15) followed by resuspension in PBS to a final concentration of 200 mg mL^−1^. NPs were characterized by dynamic light scattering (DLS) to find the NP diameter, 37 ± 4 nm.

### PNP hydrogel Preparation

Hydrogel formulations contained either 2 wt% HPMC−C_12_ and 10 wt% PEG−PLA NPs in PBS, or 1 wt% HPMC−C_12_ and 10 wt% PEG−PLA NPs in PBS. These hydrogels were made by mixing a 2:3:1 weight ratio of 6 wt% HPMC−C_12_ polymer solution, 20 wt% NP solution, and PBS containing GLP-1 RAs. The NP and aqueous components were loaded into one syringe, the HPMC-C_12_ was loaded into a second syringe and components were mixed using an elbow connector. After mixing, the elbow was replaced with a 21-gauge needle for injection.

### Rheological characterization of hydrogels

Rheological testing was performed at 25 °C using a 20-mm-diameter serrated parallel plate at a 600-μm gap on a stress-controlled TA Instruments DHR-2 rheometer. All experiments were performed at 25 °C. Frequency sweeps were performed from 0.1 to 100 rad s^−1^ with a constant oscillation strain within the linear viscoelastic regime (1%). Amplitude sweeps were performed at a constant angular frequency of 10 rad s^−1^ from 0.01% to 10000% strain with a gap height of 500 µm. Flow sweeps were performed from low to high stress with steady-state sensing. Steady shear experiments were performed by alternating between a low shear rate (0.1 s^−1^) and high shear rate (10 s^−1^) for 60 s each for three full cycles. Shear rate sweep experiments were performed from 10 to 0.001 s^−1^. Stress controlled yield stress measurements (stress sweeps) were performed from low to high stress with steady-state sensing and 10 points per decade.

### In vitro release

100 µL of each hydrogel formulation was loaded into four-inch capillaries and 400 µL of PBS medium was added slowly on top. The surrounding PBS was removed for analysis after 1, 3, 6, 12, 24, and 48 hours and at one week and two weeks after injection into the capillary, and fresh PBS was replaced after each aliquot removal. Semaglutide and liraglutide were quantified by ELISA to determine release kinetics over time.

### Animal Studies

Animal studies were performed with the approval of the Stanford Administrative Panel on Laboratory Animal Care (APLAC-32873) in accordance with NIH guidelines.

### Streptozotocin induced diabetes in rats

Male Sprague Dawley rats (Charles River) were used for experiments. Animal studies were performed in accordance with the guidelines for the care and use of laboratory animals; all protocols were approved by the Stanford Institutional Animal Care and Use Committee (Protocol #32873). Briefly, male Sprague Dawley rats 160–230 g (8–10 weeks) were weighed and fasted in the morning 6–8 h prior to treatment with nicotinamide (NA) and streptozotocin (STZ). NA was dissolved in 1X PBS and administered intraperitoneally at 110 mg/kg. STZ was diluted to 10 mg mL^−1^ in the sodium citrate buffer immediately before injection. STZ solution was injected intraperitoneally at 65 mg kg^−1^ into each rat. Rats were provided with water containing 10% sucrose for 24 h after injection with STZ. Rat BG levels were tested for hyperglycemia daily after the STZ treatment via tail vein blood collection using a hand-held Bayer Contour Next glucose monitor (Bayer). Rats were considered diabetic when at least three out of five fasting BG measurements, taken over the course of five days, fell between 130–200 mg/dL.

### 24-hour pharmacokinetics in diabetic rats

A 24-hour PK study was conducted to validate previously reported pharmacokinetic parameters (i.e., elimination half-life, volume of distribution, bioavailability, and absorption rate) of semaglutide in rats, whereby semaglutide (20 µg) was administered via bolus injection using two routes of administration, including subcutaneous (SC) and intravenous (IV). Six rats received each treatment and blood samples were collected from the tail vein every 30 minutes after treatment for eight hours, and then again at 18 hours and 24 hours.

### In-vivo pharmacokinetics and pharmacodynamics in diabetic rats

For each of the treatment groups (n = 5–6), baseline blood was collected from the tail vein at day zero and daily blood glucose measurements were taken from the tail vein using a handheld blood glucose monitor (Bayer Contour Next) for 42 days following treatment (*72*). Blood glucose was measured, immediately followed by blood samples collected from the tail vein, every day for the first seven days of the study to measure serum semaglutide concentrations using ELISA, and two times a week, thereafter. Materials for these assays were made available to us by Novo Nordisk through their compound sharing program. Plasma GLP-1 RA concentrations were measured by ELISA at each time-point and total bioavailability of semaglutide was determined at the end-point of the study. An IV glucose tolerance test (*54*) was used to group rats according to the glucose tolerance before treatment (at day −1).

### Biocompatibility

At the end of the 6-week experiment, the rats were euthanized using carbon dioxide and tissues (kidneys and liver) were collected for histology. The harvested tissue was fixed and transverse sections of the left lateral lobe and right medial lobe of the liver as well as longitudinal sections of the kidney were taken for histological analysis. Haematoxylin and eosin staining were performed by Histotec Laboratory.

### Statistical analysis

All results are expressed as a mean ± standard deviation unless specified otherwise. For in-vivo experiments, Mead’s Resource Equation was used to identify a sample size above that additional subjects will have little impact on power. Comparison between groups was conducted with the Tukey HSD test in JMP. A one-way analysis of variance (ANOVA) or a t-test was also used to compare groups. A difference of p < 0.05 was considered statistically significant.

## Acknowledgments

A.I.D. was supported by a Schmidt Science Fellows Award. C.L.M. was supported by a NSERC Postgraduate Scholarship and Stanford BioX Bowes Graduate Student Fellowship. L.T.N. was supported by a Stanford Graduate Fellowship in Science and Engineering. C.M.K. was supported by the Stanford Bio-X William and Lynda Steere Fellowship. C.K.J. and J.Y. were supported by National Science Foundation Graduate Research Fellowships. We would also like to thank all members of the Appel lab and Novo Nordisk for their useful discussion and advice throughout this project. The authors thank the Stanford Animal Diagnostic Lab and the Veterinary Service Centre staff for their technical assistance.

## Author contributions

Conceptualization: AID, EAA

Methodology: AID, CLM, LTN, EAA

Investigation: AID, CLM, LTN, KL, ANP, IAH, SWB, CMK, CKJ, JY

Visualization: AID, CLM, LTN

Funding acquisition: EAA

Project administration: EAA

Supervision: EAA

Writing – original draft: AID, EAA

Writing – review & editing: CLM, LTN, KL, ANP, IAH, SWB, CMK, CKJ, JY, LH, DBS, HBR, EAA

## Data and materials availability

All data supporting the results in this study are available within the article and its Supplementary Information. The broad range of raw datasets acquired and analyzed (or any subsets thereof), which would require contextual metadata for reuse, are available from the corresponding author upon reasonable request.

