## Supplemental Information for "Sustained Delivery of GLP-1 Receptor Agonists from Injectable Biomimetic Hydrogels Improves Treatment of Diabetes"

### Biomimetic Hydrogels for the Sustained Delivery of GLP-1 Analogues to Improve Treatment of Diabetes

**This PDF includes:**

Method S1. Method for Determining In Vitro % Cumulative Release.
Method S2. Streptozotocin/Nicotinamide (STZ/NA) induced model of type-2 diabetes in rats.
Method S3. In vivo pharmacokinetics in diabetic rats.
Method S4. Pharmacokinetic modeling.
Method S5. Method for performing an IV glucose tolerance test (IV GTT) and determining half-life.
Figure S1. Size exclusion chromatogram (SEC) trace of PEG-PLA and dynamic light scattering (DLS) of PEG-PLA nanoparticles.
Figure S2. Rheological characterization of PNP hydrogels without GLP-1 RA cargo
Figure S3. Validating Semaglutide PK parameters in rats with a 24-hour PK study
Figure S4. IV Glucose Tolerance Test (IV GTT)
Figure S5. Liver Histology – Day 42
Figure S6. Kidney Histology – Day 42
Table S1. $C_{max}$ and $C_{steady\text{-}state}$ of GLP-1 RA Hydrogel Formulations
Table S2. P-Values for $C_{max}$
Table S3. P-Values for $C_{steady\text{-}state}$
Table S4. P-Values for Blood Glucose
Table S5. P-Values for Weight
Table S6. Liraglutide Rat Modeling Parameters
Table S7. Liraglutide Human Modeling Parameters

**Method S1. Method for determining in vitro % cumulative release.**
In vitro cumulative release was determined by taking the ratio of $\frac{M_t}{M_{inf}}$, where $M_t$ is the amount of drug released at time *t*, and $M_{inf}$ is the total amount of drug within the gel. Quantity of drug was determined using either a Semaglutide- or Liraglutide-specific ELISA kit (BMA Biomedical).

**Method S2. Streptozotocin/Nicotinamide (STZ/NA) induced model of type-2 diabetes in rats.**
Male Sprague Dawley rats (Charles River) were used for experiments. Animal studies were performed in accordance with the guidelines for the care and use of laboratory animals; all protocols were approved by the Stanford Institutional Animal Care and Use Committee. The protocol used for STZ/NA induction was adapted from the protocol by Ting Chen, Leonid Kagan and Donald E. Mager (*1*). Briefly, male Sprague Dawley rats 160–230 g (8–10 weeks) were weighed and fasted 6–8 hours prior to treatment with STZ/NA. Rats were injected IP with 110 mg/kg nicotinamide. Fifteen minutes later, rats were treated with STZ. STZ was diluted to 10 mg/mL in the sodium citrate buffer immediately before injection. STZ solution was injected intraperitoneally at 65 mg/kg into each rat. Rats were provided with water containing 10% sucrose for 24 hours after injection with STZ/NA. Rat blood glucose levels were tested for hyperglycemia daily after the STZ/NA treatment via a tail vein blood collection using a handheld Bayer Contour Next glucose monitor (Bayer). Diabetes was defined as having three consecutive blood glucose measurements ranging between 130–200 mg/dL in non-fasted rats.

**Method S3. In vivo pharmacokinetics in diabetic rats and PK modeling from a gel.**
A 24-hour PK study was conducted to validate the PK parameters of Semaglutide in rats, as well as to model PK from our PNP hydrogel. To do this we need to know the elimination half-life (clearance rate) of Semaglutide which can be calculated by fitting an exponential curve to the IV PK data (one compartment model). We also want to know the absorption rate of Semaglutide from the SC space (two compartment model). We can use the elimination half-life from the IV data to define the clearance rate from compartment two (blood) and solve for the absorption rate from compartment one into compartment two (SC into blood). Both rates provide sufficient information to then project what long-term release from the gel will look like based on diffusion of cargo from the gel.

To conduct the 24-hour PK study, diabetic rats were first fasted for 4-6 hours. Semaglutide was administered intravenously as a soluble solution in PBS buffer (20 µg). 80 µL of blood was collected at 0 min, 30 min, 60 min, 90 min, 120 min, 150 min, 180 min, 210 min, 240 min, 270 min, 300 min, 330 min, 360 min, 390 min, 420 min, 450 min, 480 min, 18 h and 24 h after injection. Samples were analyzed using a Semaglutide-specific ELISA purchased from BMA Biomedical.

**Method S4. Pharmacokinetic modeling.**
The differential equations and the analytical solution for a standard one compartment pharmacokinetic model with first order reaction kinetics are described in Zou, et al (*2*). Briefly, the form of the analytical solution of the single compartment model for drug serum concentration as a function of time, $C(t)$, used in our modeling is as follows:

$$C(t) = \frac{F k_{abs} M_0}{V_d(k_{abs} - k_{elim})}\left(e^{-k_{elim}} - e^{-k_{abs} t}\right) \quad \text{(S1)}$$

where $F$ = bioavailability, $k_{abs}$ = rate constant of absorbance from subcutaneous space into the bloodstream, $M_0$ = initial dose of drug, $V_d$ = volume of distribution, $k_{elim}$ = rate constant of drug elimination, and $t$ = time.

The following differential equations and conditions were used to describe mass transport. $M_1$ refers to drug mass in the subcutaneous hydrogel depot compartment, $M_2$ refers to drug mass in the subcutaneous compartment, $M_3$ refers drug mass in the blood serum compartment and $k_{dpt}$ is the rate constant of drug release from the hydrogel depot.

$$\frac{dM_1(t)}{dt} = -k_{dpt}M_1(t) \qquad M_1(0) = M_0 \tag{S2}$$

$$\frac{dM_2(t)}{dt} = k_{dpt}M_1(t) - k_{abs}M_2(t) \qquad M_2(0) = 0 \tag{S3}$$

$$\frac{dM_3(t)}{dt} = k_{abs}FM_2(t) - k_{elim}M_3(t) \qquad M_3(0) = 0 \tag{S4}$$

$$C(t) = \frac{M_3(t)}{V_d} \tag{S5}$$

These equations were solved for the following analytical solution:

$$C(t) = \frac{Fk_{dpt}M_0k_{abs}e^{-k_{elim}t}}{V_d(k_{dpt}-k_{abs})(k_{elim}-k_{dpt})(k_{elim}-k_{abs})}\left[(k_{elim} - k_{dpt})e^{(k_{elim}- k_{abs})t} + (k_{abs} - k_{elim})e^{(k_{elim}- k_{dpt})t} + k_{dpt} - k_{abs}\right] \tag{S6}$$

**Method S5. Method for performing an IV glucose tolerance test (IV GTT).**

An IV glucose tolerance test (IV GTT) was used as a metric to assess insulin release and insulin resistance at baseline before treatment (at day -1) and at the end of the study (day 42), according to previously reported procedures (*1*). An intravenous glucose tolerance test (IV GTT) was performed for all groups after an overnight fast. A 50 wt% solution of glucose was intravenously injected at 2.78 mmol/kg, and blood glucose was measured at –60, –20, –10, 5, 10, 15, 20, 30, 45, 60, and 120 min. We paired rats with similar glucose tolerance into bins and then treatment groups were randomized to each rat.

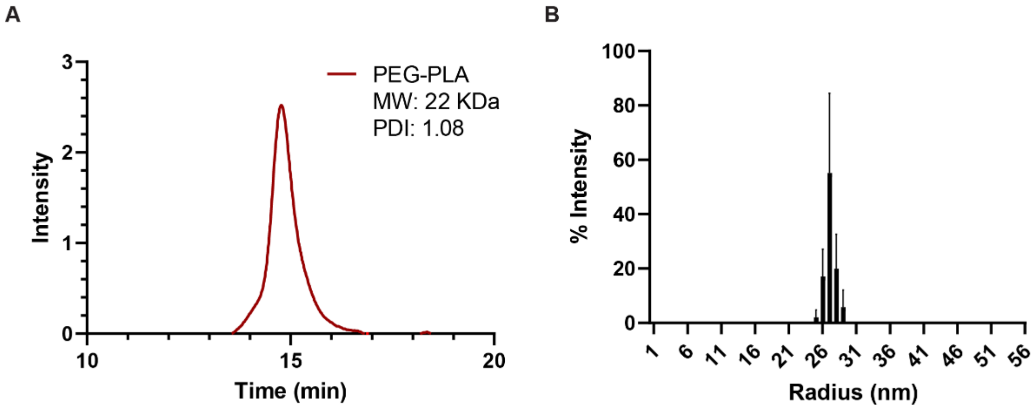


**Figure S1. Size exclusion chromatogram (SEC) trace of PEG-PLA and dynamic light scattering (DLS) of PEG-PLA nanoparticles.**

A) Gel permeation chromatography characterization of the PEG-PLA polymer showing a single peak corresponding to the PEG-PLA block co-polymer ($M_n$ = 22 kDa; Đ = 1.08). B) DLS characterization of the PEG-PLA nanoparticles after precipitation ($D_H$ = 33.2, PDI = 0.038).

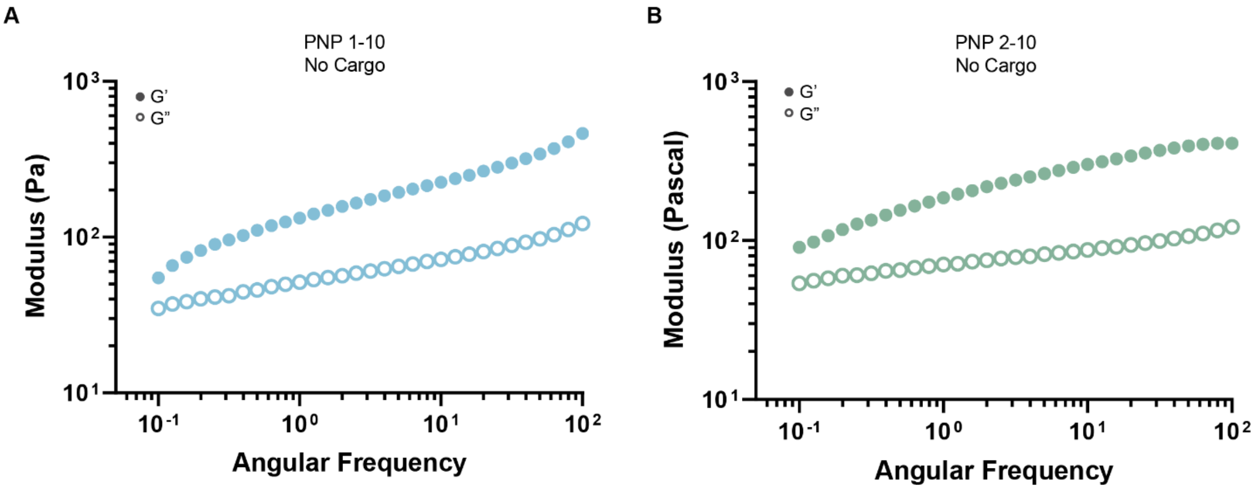


**Figure S2. Rheological characterization of PNP hydrogels without GLP-1 RA cargo.** Rheology of the A) PNP-1-10 and B) PNP-2-10 hydrogel control formulations without GLP-1 RAs. Frequency sweep (% strain = 1%) for all formulations at 25 °C.

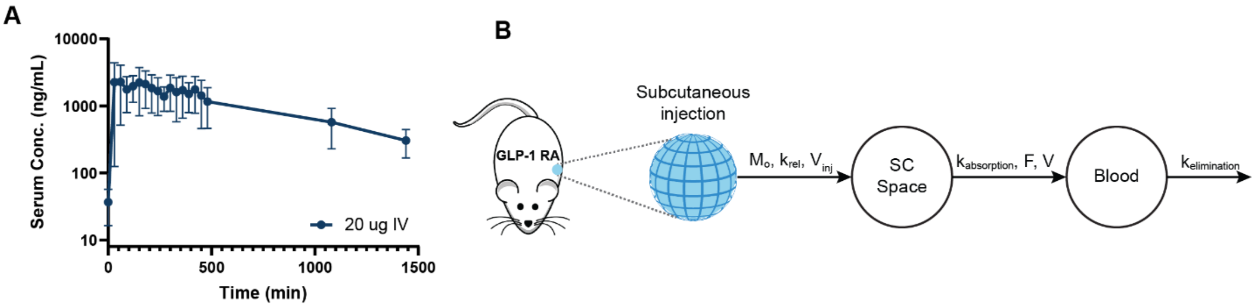


**Figure S3. Validating Semaglutide PK parameters in rats with a 24-hour PK study.** Semaglutide pharmacokinetics in a T2D rat model as determined by ELISA. A) PK profile for Semaglutide administered intravenously as a soluble solution in PBS buffer (20 µg). B) Two-compartment model describing the mass transport of cargo from a subcutaneous injection.

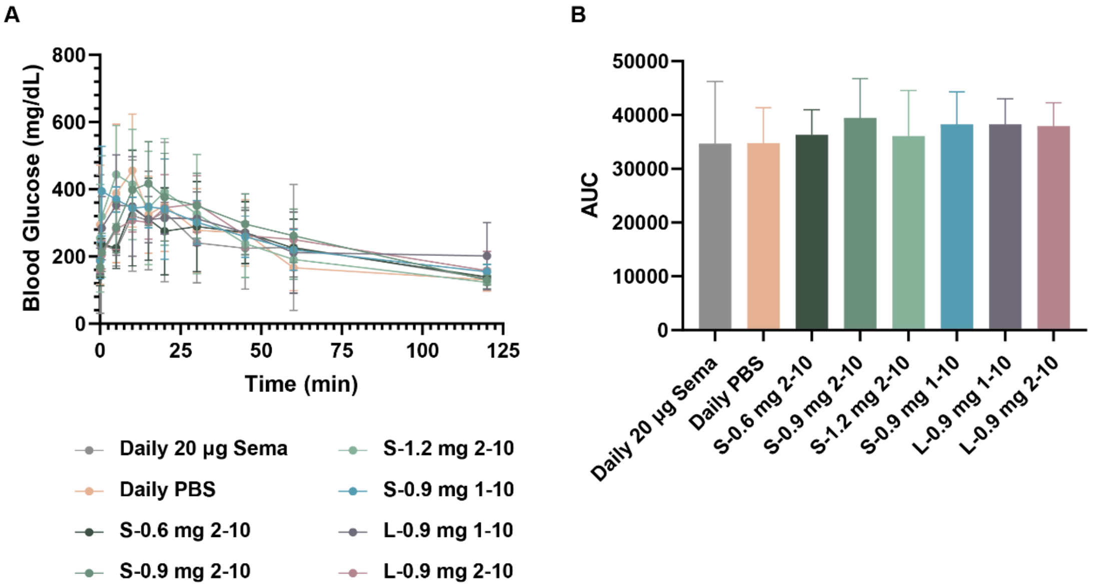


**Figure S4. IV Glucose Tolerance Test (IV GTT).**
An IV glucose tolerance test (IV GTT) was conducted to group the rats into treatment groups. A total of eight treatment groups were evaluated. A) Blood glucose was measured at –60, –20, –10, 5, 10, 15, 20, 30, 45, 60, and 120 min. B) Using the area under the curve (AUC), rats with similar glucose tolerance were paired and then randomized into treatment groups.

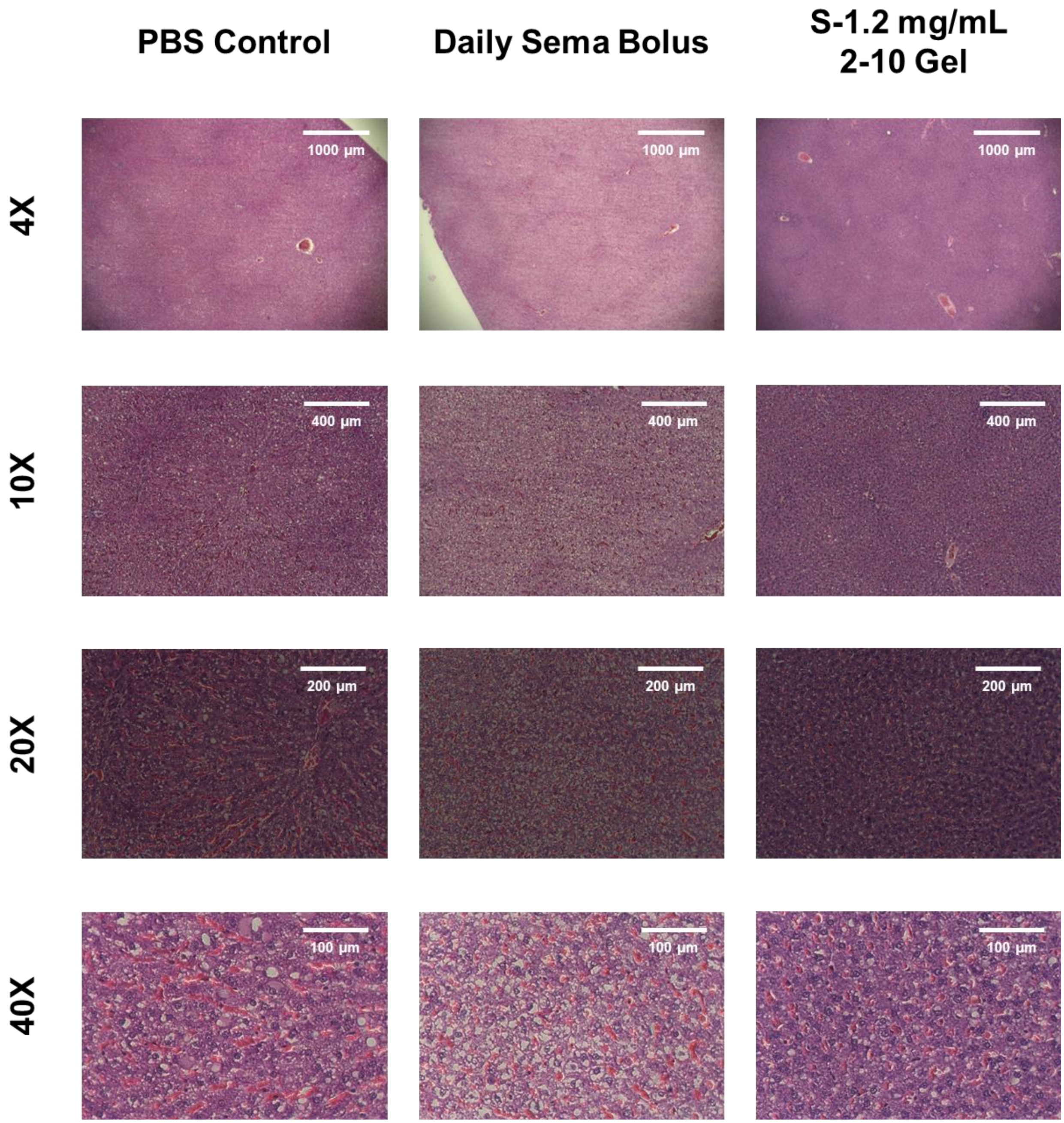


**Figure S5. Liver Histology – Day 42.**
Hematoxylin and eosin (H&E) staining, representative images at Day 42. Images show cross sections of liver after 42 days.

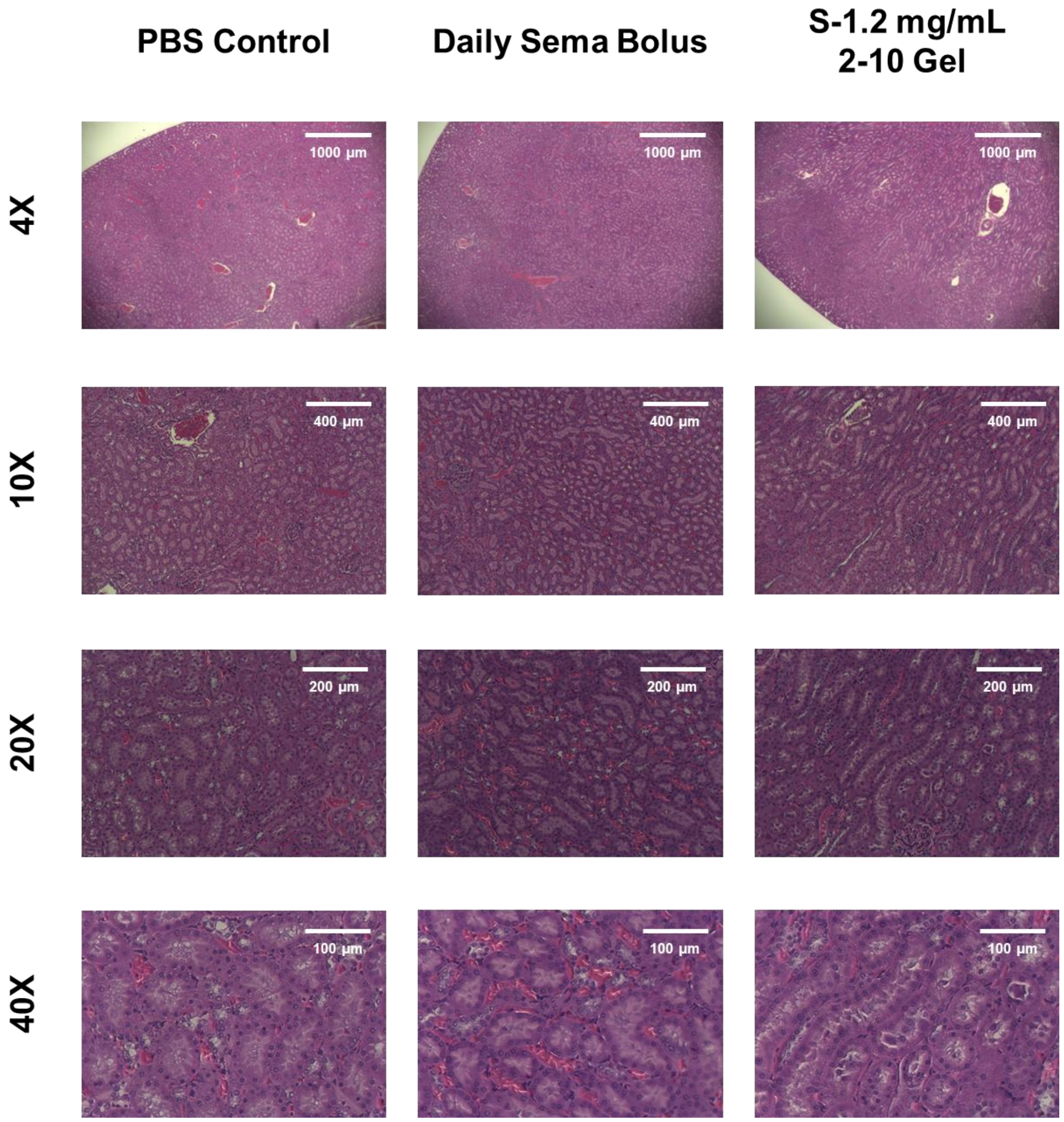


**Figure S6. Kidney Histology – Day 42.**
Hematoxylin and eosin (H&E) staining, representative images at Day 42. Images show cross sections of kidney after 42 days.

**Table S1. Pharmacokinetic Parameters of GLP-1 RAs in Rats**

| GLP-1 RA | Elimination half-life |
|---|---|
| Semaglutide | 7 hr (*3*) |
| Liraglutide | 4 hr (*4*) |

**Table S2. $C_{max}$ and $C_{steady\text{-}state}$ of GLP-1 RA Hydrogel Formulations**

| Formulation | Time to $C_{max}$ | $C_{max}$ (ng/mL) | Time to $C_{steady\text{-}state}$ | $C_{steady\text{-}state}$ (ng/mL) |
|---|---|---|---|---|
| Sema Bolus | Day 34 | 280 ± 60 | Day 7 | 170 ± 60 |
| S-0.6 mg 2-10 | Day 1 | 1110 ± 430 | Day 7 | 66 ± 50 |
| S-0.9 mg 2-10 | Day 1 | 3800 ± 900 | Day 7 | 60 ± 40 |
| S-1.2 mg 2-10 | Day 1 | 4400 ± 1200 | Day 7 | 75 ± 10 |
| S-0.9 mg 1-10 | Day 1 | 5900 ± 2100 | Day 7 | 40 ± 40 |
| L-0.9 mg 1-10 | Day 9 | 620 ± 110 | Day 1 | 270 ± 190 |
| L-0.9 mg 2-10 | Day 20 | 930 ± 350 | Day 1 | 410 ± 200 |

**Table S3. P-Values for $C_{max}$**

| Formulation Comparison | $C_{max}$ P-value |
|---|---|
| S-0.6 mg 2-10 vs. S-1.2 mg 2-10 | 0.0003* |
| S-0.6 mg 2-10 vs. S-0.9 mg 2-10 | 0.0010* |
| S-1.2 mg 2-10 vs. S-0.9 mg 2-10 | 0.6000* |
| S-0.9 mg 2-10 vs. S-0.9 mg 1-10 | 0.0700** |
| L-0.9 mg 2-10 vs. L-0.9 mg 1-10 | 0.0900** |

*P-value determine using ordinary one-way ANOVA
**P-value determine using unpaired t-test

**Table S4. P-Values for $C_{steady\text{-}state}$**

| Formulation Comparison | $C_{steady\text{-}state}$ P-value |
|---|---|
| S-0.6 mg 2-10 vs. S-1.2 mg 2-10 | 0.8* |
| S-0.6 mg 2-10 vs. S-0.9 mg 2-10 | 0.6* |
| S-1.2 mg 2-10 vs. S-0.9 mg 2-10 | 0.3* |
| S-0.9 mg 2-10 vs. S-0.9 mg 1-10 | 0.3** |
| L-0.9 mg 2-10 vs. L-0.9 mg 1-10 | 0.02** |

*P-value determine using ordinary one-way ANOVA
**P-value determine using unpaired t-test

**Table S5. P-Values for Blood Glucose**

| Formulation Comparison | P-value |
|---|---|
| PBS Bolus vs. S-0.6 mg 2-10 | 0.0030 |
| PBS Bolus vs. S-0.9 mg 2-10 | 0.0139 |
| PBS Bolus vs. S-1.2 mg 2-10 | <0.0001 |
| PBS Bolus vs. S-0.9 mg 1-10 | <0.0001 |
| PBS Bolus vs. L-0.9 mg 2-10 | 0.0004 |
| PBS Bolus vs. L-0.9 mg 1-10 | 0.0006 |

P-values determined using unpaired t-test

**Table S6. P-Values for Weight**

| Formulation Comparison | P-value |
|---|---|
| PBS Bolus vs. S-0.6 mg 2-10 | 0.0792 |
| PBS Bolus vs. S-0.9 mg 2-10 | 0.0040 |
| PBS Bolus vs. S-1.2 mg 2-10 | 0.0061 |
| PBS Bolus vs. S-0.9 mg 1-10 | 0.1824 |
| PBS Bolus vs. L-0.9 mg 2-10 | 0.3256 |
| PBS Bolus vs. L-0.9 mg 1-10 | 0.0265 |

P-values determined using unpaired t-test

**Table S7. Liraglutide Rat Modeling Parameters**

| Parameter | Gel | PBS |
|---|---|---|
| Bioavailability | 0.82 | 0.82 |
| Injection volume | 500 µL | 500 µL |
| Concentration | 2.4 mg/mL | 1.8 mg/mL |
| Mass of drug | 1,200 µg | 900 µg |
| Blood volume | 25 mL | 25 mL |
| Release rate (k) | 0.02 $day^{-1}$ | 0.02 $day^{-1}$ |
| Release coefficient (n) | 0.7 | 0.7 |
| $t_{1/2 \text{ in blood}}$ | 0.167 day | 0.167 day |
| $t_{1/2 \text{ in SC}}$ | 0.9504 min | 0.9504 min |

**Table S8. Liraglutide Human Modeling Parameters**

| Parameter | Gel | PBS |
|---|---|---|
| Bioavailability | 0.84 | 0.84 |
| Injection volume | 1,250 µL | 1,250 µL |
| Concentration | 20 mg/mL | 1 mg/mL |
| Mass of drug | 20,000 µg | 1,000 µg |
| Blood volume | 5,000 mL | 5,000 mL |
| Release rate (k) | 0.08 $day^{-1}$ | 0.08 $day^{-1}$ |
| Release coefficient (n) | 1 | 1 |
| $t_{1/2 \text{ in blood}}$ | 0.5417 day | 0.5417 day |
| $t_{1/2 \text{ in SC}}$ | 0.25 day | 0.25 day |
